# A maladaptive combination of traits contributes to the maintenance of a stable hybrid zone between two divergent species of *Drosophila*

**DOI:** 10.1101/138388

**Authors:** Brandon S. Cooper, Alisa Sedghifar, W. Thurston Nash, Aaron A. Comeault, Daniel R. Matute

**Author notes:** Correspondence: Division of Biological Sciences, University of Montana, Missoula, 32 Campus Drive, Missoula, MT 59812, USA. Biology Department, University of North Carolina, Chapel Hill, 250 Bell Tower Drive, Genome Sciences Building, Chapel Hill, NC 27510, USA.

## Abstract

Geographical areas where two species come into contact and hybridize serve as natural laboratories for assessing mechanisms that limit gene flow between species. The ranges of about half of all closely related *Drosophila* species overlap, and the genomes of several pairs reveal signatures of past introgression. However, only two contemporary hybrid zones have been characterized in the genus, and both are recently diverged sister species (*D. simulans-D. sechellia*, Ks = 0.05; *D. yakuba-D. santomea*, Ks = 0.048). Here we present evidence of a new hybrid zone, and the ecological mechanisms that maintain it, between two highly divergent *Drosophila* species (Ks = 0.11). On the island of Bioko in west Africa, *D. teissieri* occupies mostly forests, *D. yakuba* occupies mostly open agricultural areas, and recently, we discovered that hybrids between these species occur near the interface of these habitats. Genome sequencing revealed that all field-sampled hybrids are F_1_ progeny of *D. yakuba* females and *D. teissieri* males. We found no evidence for either advanced-generation hybrids or F_1_ hybrids produced by *D. teissieri* females and *D.yakuba* males. The lack of advanced-generation hybrids on Bioko is consistent with mark-recapture and laboratory experiments that we conducted, which indicate hybrids have a maladaptive combination of traits. Like *D. yakuba*, hybrids behaviorally prefer open habitat that is relatively warm and dry, but like *D. teissieri*, hybrids have low desiccation tolerance, which we predict leaves them physiologically ill-equipped to cope with their preferred habitat. These observations are consistent with recent findings of limited introgression in the *D. yakuba* clade and identify an ecological mechanism for limiting gene flow between *D. yakuba* and *D. teissieri*; namely, selection against hybrids that we have documented, in combination with hybrid male sterility, contributes to the maintenance of this narrow (~30m), stable hybrid zone centered on the forest-open habitat ecotone. Our results show how a deleterious combination of parental traits can result in unfit or maladapted hybrids.

## INTRODUCTION

Genetically distinct populations of closely related species often come into contact, mate, and produce offspring in narrow zones of hybridization (Barton and Hewitt 1985; Hewitt 1988). These geographical areas offer windows on the evolutionary processes that lead to the origin and maintenance of species (Harrison 1990), and have been of interest to evolutionary biologists for over a century (Harrison and Larson 2016). Incipient species could potentially fuse into a single entity if fertile hybrids are produced and successfully backcross into one or both parental species, but because selection has not tested the combinations of alleles found in divergent species, hybrids are often less fit than either parental species (Dobzhansky and Dobzhansky 1937; Muller 1942; Turelli et al. 2001). For example, hybrids have been shown to have both developmental (Mendelson 2003; Coyne and Orr 2004; Moyle et al. 2004; Maheshwari and Barbash 2011) and behavioral defects (Coyne and Orr 1989; Funk et al. 2006; Turissini et al. 2017). These reproductive isolating mechanisms should contribute to the maintenance of species in areas of contact if they prevent the formation of hybrids, or if they lead to the production of sterile and/or low fitness hybrids.

Estimates from published data indicate that hybridization is common (Stebbins 1959; Barton and Hewitt 1985). For example, approximately 25% of vascular plant species and 10% of animal species show evidence of past hybridization (Grant and Grant 1992; Mallet 2005; Mallet et al. 2016). In the laboratory, there is some hybrid viability after divergence times of 60 million years (my) in birds (Price and Bouvier 2002), 30 my in plants (Widmer et al. 2009), and 15 my in *Drosophila* (Turissini and Matute in revision). In the most extreme cases, successful hybridization seems possible between species pairs of lancelets (120 my diverged) (Holland et al. 2015), ferns (60 my diverged) (Rothfels et al. 2015), and fungi (at least 3% nuclear divergence) (Stukenbrock et al. 2012). Thus, while estimated divergence times remain error prone and inexact, these data indicate that hybridization between highly diverged species is at least possible.

About half of all closely related *Drosophila* species have overlapping geographical ranges (Turelli et al. 2014), and several species pairs show evidence of past introgression (Shoemaker et al. 1999; Jaenike et al. 2006; Kulathinal et al. 2009; Garrigan et al. 2012; Brand et al. 2013; Lohse et al. 2015), but only two contemporary hybrid zones have been identified and well described in the genus. Both hybrid zones involve closely related sister species in the *D. melanogaster* subgroup: *Drosophila simulans* and *D. sechellia* (Ks = 0.05) hybridize in the central islands of the Seychelles archipelago where human populations have facilitated the spread of *D. simulans* into the ancestral range of *D. sechellia* (Matute and Ayroles 2014), and *D. yakuba* hybridizes with the endemic species *D. santomea* (Ks = 0.048) on the island of São Tomé in west Africa (Lachaise et al. 2000; Llopart et al. 2005a). Along the altitudinal transect of Pico de São Tomé, *D. yakuba* occurs at low elevations (below 1,450 m), *D. santomea* occurs at high elevations (1,150 m and 1,800 m), and where their ranges overlap 3% of the sampled *D. yakuba* clade individuals are hybrids (Llopart et al. 2005b; Comeault et al. 2016). Past evidence suggested extensive mitochondrial (mt) introgression between these species (Bachtrog et al. 2006; Llopart et al. 2014), but more recent analyses of whole mt, *Wolbachia*, and nuclear genomes suggest less introgression (Turissini and Matute in revision; Turelli, Conner, Turissini, Matute, and Cooper unpublished).

*Drosophila teissieri* diverged from *D. yakuba* and *D. santomea* approximately ~3.0 my ago (Turissini and Matute in revision, Turelli, Conner, Turissini, Matute, and Cooper unpublished) and is thought to share a large ancestral range with *D. yakuba* across much of the African continent (Lachaise et al. 1981; Cobb et al. 2000). Due to widespread deforestation *D. teissieri* is now distributed across fragmented tropical forests (mostly at altitudes over 500 m above sea level) in continental Africa and in west Africa on the island of Bioko (Lachaise et al. 1981; Cobb et al. 2000). In contrast, *D. yakuba* is a human commensal and occurs in disturbed, open habitats across the African continent and on Bioko, in addition to São Tomé and other islands (Matute 2010; Yassin et al. 2016; Yassin 2017). *Drosophila teissieri* is thought to be associated with *Parinari* fruits, which may also restrict its current geographic range (Rio et al. 1983; David et al. 2007; Comeault et al. in press). *Drosophila teissieri* can be hybridized with *D. yakuba* under laboratory conditions, but matings are rare (~9% of pairs mate, Turissini et al. 2015).Nevertheless, and despite being highly diverged (Ks = 0.112, Turissini et al. 2015), reciprocal *D. teissieri-D. yakuba* hybrid females are fertile, providing the possibility for gene exchange between species. In contrast, hybrid males are sterile. Hybrids from other *Drosophila* species pairs with a similar level of divergence are inviable or sterile (e.g., *D. melanogaster-D. simulans* hybrids, Ks = 0.10; Matute et al. 2010), which makes *D. yakuba* and *D. teissieri* one of the most diverged *Drosophila* pairs to produce fertile progeny in the laboratory. The geographical overlap of *D. teissieri* and *D. yakuba* throughout areas of Africa, combined with laboratory crossing results and genomic observations of some introgression, suggests contemporary hybridization between these species is possible.

Here, we report a new *Drosophila* hybrid zone that we discovered by intensely sampling Bioko, which is composed of primary forest, open habitat devoted to subsistence agriculture, and secondary forest that has grown in the last 50 years. We present multiple lines of evidence indicating that *D. yakuba* and *D. teissieri* hybridize at the interface of secondary forests (preferred by *D. teissieri*) and open habitats (preferred by *D. yakuba*). Sampling of hybrids in both 2009 and 2013 suggests some stability of this zone, and genome sequencing reveals that all sampled hybrids are F_1_ progeny of *D. yakuba* females and *D. teissieri* males. Field and laboratory experiments indicate that hybrids have *D. yakuba’s* behavioral preference for warm and dry conditions, and *D. teissieri’s* limited physiological tolerance for low humidity conditions—a seemingly maladaptive combination of parental traits. We predict that this maladaptive combination of parental traits, in combination with hybrid male sterility, maintains this narrow (~30 m), stable hybrid zone centered on the forest-open habitat ecotone.

## METHODS

### Distinguishing among *D. yakuba, D. teissieri*, and F_1_ hybrids using morphology

We sought to heavily sample Bioko for *D. yakuba, D. teissieri*, and their putative hybrids. However, conducting experiments in the field requires a method for reliably identifying and distinguishing living individuals of each genotype. Based on our experience with *D. yakuba* clade flies, we predicted that three male morphological characteristics would enable us to achieve this goal: the number of chitinized spines on anal plates, the number of sex combs on forelegs, and the lengths of tibia. We first measured each trait in a training set of *D. yakuba* (*N*=500) and *D. teissieri* (*N*=500) males, and also in F_1_(♀*tei* × ♂ *yak*) (*N* = 500) and F_1_(♀*yak* × ♂ *tei*) (*N* = 500) hybrid males that we created in the laboratory—F_0_ parental crosses are listed in parentheses for each of the two F_1_ genotypes. This provided a distribution of values for each trait within each genotype. We next blindly measured each of the three morphological traits in an additional set of 100 males of each genotype and calculated Mahalanobis distances for each individual (Mahalanobis 1936). The Mahalanobis distance between a focal individual (*i*) and the average for a given genotype (i.e., centroids for *D. teissieri, D. yakuba*, and the two hybrid F_1_ genotypes estimated using the training set) was calculated as:

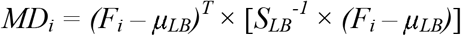

where the super-index T denotes matrix transpose, S denotes the covariance matrix of a given dataset, *F_i_* is the vector of phenotypic observations in a focal individual, *i*, and *μ_LB_* is the vector of average phenotypic observations of the training set. To estimate the accuracy of this approach, we calculated the proportion of our blind assignments that were correct. Mahalanobis distances were later used in the field to distinguish genotypes, and the field assignments were later verified using genomic sequencing (see below).

### Geographic distributions of *D. yakuba, D. teissieri*, and their hybrids on Bioko

To determine the geographical distributions of *D. yakuba* and *D. teissieri*, and to identify any evidence of hybridization between these species, we carried out collection expeditions in July 2009 and September 2013 on Bioko. The 2009 sampling was conducted over a period of 25 days and involved only two altitudes (1,200 and 1,650 m). In 2013, we widely sampled the north portion of the island at seven altitudes (200, 650, 1,200, 1,420, 1,650, 1,850, and 2,020 m) that covered approximately 20 km. In both years we collected flies using bottle traps containing fermented banana mash that were hung from trees with tape at a height of 1.2 m to 1.5 m from the ground. In 2013, we collected over a period of 14 days when all of the traps were in place for each sampled altitude. On day 15, the traps at 2,020 m became inaccessible for safety reasons. Every three days, we replaced the bait in each trap. Flies were aspirated from traps by mouth (1135A Aspirator – BioQuip; Rancho Domingo, CA) every 24 hr and transferred to empty glass vials with wet paper balls (to provide humidity) where they remained for a period of up to three hours. Flies were then lightly anesthetized using FlyNap (Carolina Biological Supply Company, Burlington, NC) and sorted by species using Mahalanobis distances. We focused on taxonomic identification of only males since the identification of living females is not possible (Markow and O’Grady 2005). While our traps attracted other species known to occur on Bioko, our analyses were restricted to *D. yakuba* clade males. The flies sampled in 2013 were housed in 10 mL plastic containers that contained instant fly food (Carolina Biological, Burlington, NC) supplemented with boiled banana paste for up to 10 days. These flies were then used to assess the habitat preferences of *D. yakuba, D. teissieri*, and their hybrids (see below).

We also sampled flies from the surfaces of ripe mangoes, papayas, figs, and *Parinari* fruits in forests using aspirators; and we netted flies off of vegetable debris in the main population centers of Malabo, Riaba, Luba, and Moka. Single females were placed in 10 mL plastic containers that contained instant fly food (Carolina Biological, Burlington, NC) supplemented with boiled banana paste. Once in the laboratory, F_1_ male morphology was evaluated to identify species. All stocks and populations were reared on standard *Drosophila* cornmeal medium at 24°C under a 12 hr light/dark cycle.

### Genome sequencing of naturally sampled individuals

We sequenced a subset of the male individuals sampled from Bioko in 2013 to confirm our genotype assignments based on Mahalanobis distances in the field. Genomic DNA was extracted from each individual using the Beckman-Coulter DNAdvance magnetic bead protocol for insects (Beckman Coulter, Indianapolis, IN, USA). Libraries were made for each individual using the Tagmentase protocol detailed in Picelli et al. (2014), and barcoded for multiplexing (see supplement for barcode sequences). Libraries were then sequenced to low coverage with Illumina 100 bp single end reads (Cornell Genomics Facility). In total, 20 parental *D. teissieri*, 17 parental *D. yakuba*, and 19 individuals of putative hybrid phenotype were sequenced to sufficient coverage (i.e., > 10,000 reads). The mean number of markers per individual was 6.869, and the mean coverage was 2.29X.

Genotypes of the individuals were determined by the Multiplexed Shotgun Genotyping (MSG) pipeline described by Andolfatto et al. (2011), which uses a hidden Markov model (HMM) to assign ancestry along a genome with low-coverage read data. Because this approach utilizes linkage disequilibrium on a large physical scale, it is well-suited for assigning genome-wide ancestry in recently formed hybrids (within several generations of backcrossing). The *D. yakuba* Flybase assembly (version 1.05) was repeat-masked (Smit et al. 2013-2015) and used as the first parental reference. The second parental input for MSG was made by mapping reads from the outbred individual Bioko_cascade_2_2 (Turissini and Matute in revision) to the *D. yakuba* flybase assembly and creating an updated FASTA file with genotype calls from GATK (McKenna et al. 2010). The updated FASTA file uses only single nucleotide polymorphisms (SNPs), and masks all inferred indels plus 5 bases both up and downstream. Some regions of the genome are error-prone in terms of assigning ancestry, due in part to low sequence divergence, poor reference genome assembly, or high polymorphism in the parental species. To reduce the rate of miscalled ancestry, known intermediate-frequency SNPs in each parental population (identified by realigning whole-genome Illumina sequencing data to the *D. yakuba* reference) were masked in the corresponding reference genomes using sequence data from several wild-caught individuals: Bioko_Cascade_21 (*yak*), Bioko_Cascade_19_16 (*yak*), Bioko_NE_4_6 (*yak*), Bioko_Balancha_1 (*tei*), Bioko_cascade_4_3 (*tei*), Bioko_House_Bioko (*tei*), Bioko_cascade_4_2 (*tei*), Bioko_cascade_4_1 (*tei*), Bioko_cascade_2_4 (*tei*), Bioko_cascade_2_2 (*tei*), Bioko_cascade_2_1 (*tei*). The performance of MSG is also influenced by user-specified parameters in the HMM, in particular, those that describe the error-rate of genotypes and the rate at which transitions between ancestries occur (*rfac, deltapar1*, and *deltapar2*). To determine the appropriate parameter values for this study, MSG was run iteratively on pure-species individuals while adjusting parameters to achieve the lowest genotyping error rate, assuming that all individuals are homozygous genome-wide for their ancestry. After parameter-tuning, MSG was run on all individuals and ancestry-genotypes were called with a posterior probability filter of 0.99.

### Mark-recapture experiments in the field

We sought to determine the habitat preferences (forest vs. open habitat) of *D. teissieri, D. yakuba*, and their hybrids. In 2013, we completed a preliminary experiment to first determine the distance flies travel in ~24 hr. *Drosophila melanogaster* subgroup species males (*D. yakuba, D. teissieri, D. simulans*, and *D. melanogaster*) were collected using baited traps (described above), anesthetized with FlyNap, and assigned to one of the four species based on genital morphology (Markow and O’Grady 2005). This approach yielded at least 500 male individuals of each species that we dusted with micronized fluorescent powder (Signal green; Day-Glo Color Corporation, Cleveland, OH), and released at once (*N* = 2,000 released) from a single location (1,650 m). Prior to this release, traps that consisted of small buckets with a mixture of banana and yeast were placed every 100 m in six radially arranged transects around the release area (8 traps per transect for a total of 49 traps, including a trap at the intersection of the transects). Traps were sampled between 23 and 25 hr later, and the number of recaptured flies was determined using a UV light. Species were again identified by their genital morphology. These experiments revealed that *D. yakuba* and *D. teissieri* flies rarely moved more than 100 m over a period of 24 hr (Table S1).

We determined habitat preferences of *D. teissieri* (*N* = 1,076), *D. yakuba* (*N* = 1,211), and hybrid males (*N* = 144) sampled across the altitudinal transect in 2013 (Table 1). This experiment was completed in independent blocks (*N* = 3). Flies were lightly dusted with one of three types of micronized fluorescent powder (Signal green, Horizon blue, and Fire orange; Day-Glo Color Corporation, Cleveland, OH) and allowed to spread from a trap placed at the center of the forest-open habitat ecotone, at an altitude of approximately 1,650 m. After an average of 24 hr, we recaptured flies by netting over traps that consisted of small buckets with a mixture of banana and yeast. Buckets were evenly spaced by ~10 m, with 11 running into the forest and 11 into the open habitat perpendicular to the interface of the two habitats. This design covered a total length of more than 100 m from the center of the ecotone, which our preliminary analysis suggested is sufficient (Table S1). Upon collection, genotypes were identified by dust color under UV light, and the dust-color was changed for each block such that each genotype experienced each color once. Testes were dissected from all recaptured males, mounted on slides in Ringer’s solution, and evaluated for sperm motility. Hybrid males are sterile and have no motile sperm, while non-hybrids are fertile with motile sperm (Turissini et al. 2015).

**TABLE 1.**
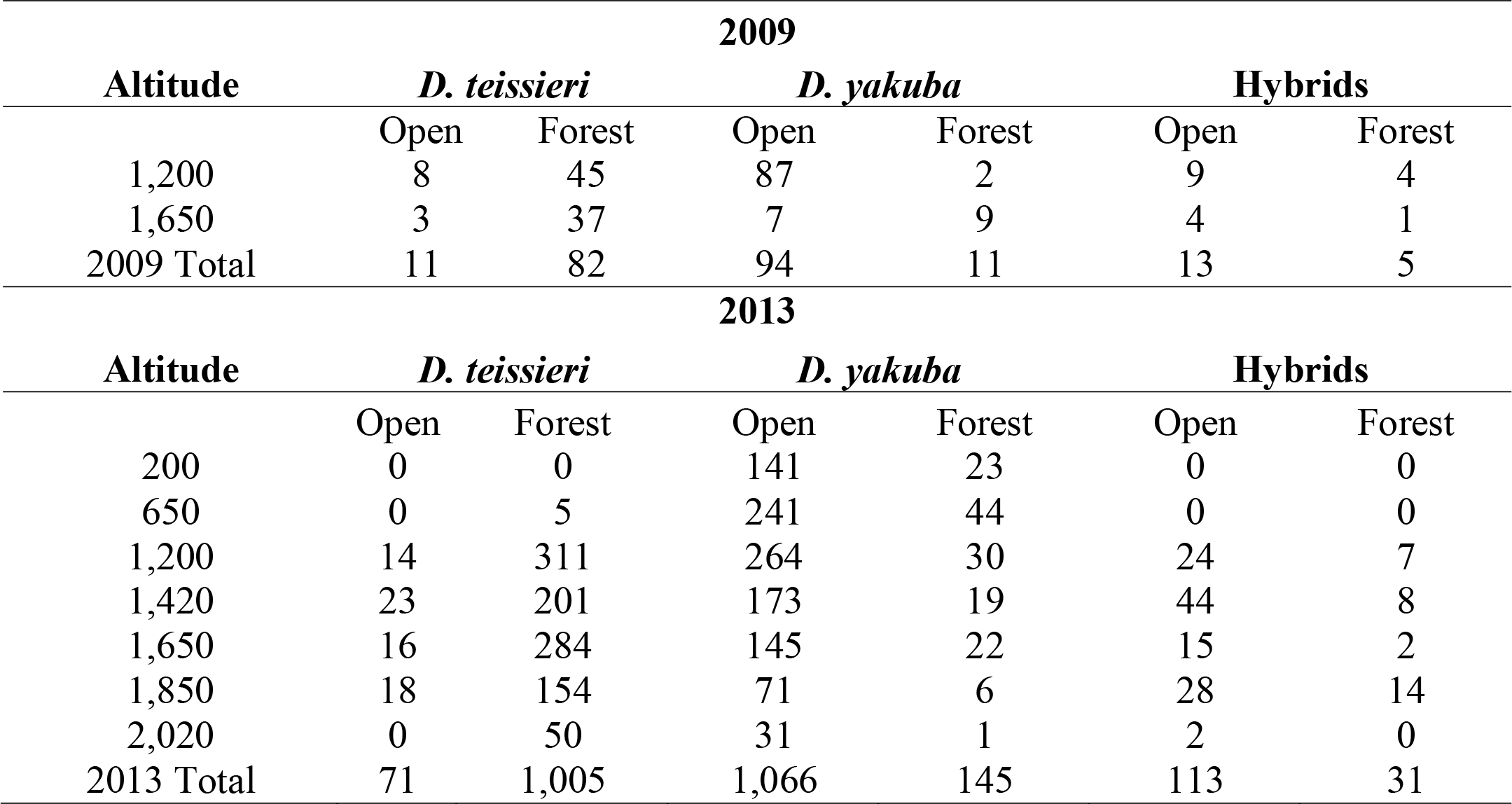
The distributions of *D. yakuba, D. teissieri*, and their hybrids in open and forest habitats, at different altitudes. Flies sampled in 2013 were used in the mark-recapture release experiment, and only flies that survived until this experiment are included here. While sampling of each habitat is reported for hybrids, these genotypes were circumscribed to the center of the forest-open habitat ecotone.

| **2009** |  |  |  |  |  |  |
| --- | --- | --- | --- | --- | --- | --- |
| **Altitude** | ***D. teissieri*** |  | ***D. yakuba*** |  | **Hybrids** |  |
|  | Open | Forest | Open | Forest | Open | Forest |
| 1,200 | 8 | 45 | 87 | 2 | 9 | 4 |
| 1,650 | 3 | 37 | 7 | 9 | 4 | 1 |
| 2009 Total | 11 | 82 | 94 | 11 | 13 | 5 |
| **2013** |  |  |  |  |  |  |
| **Altitude** | ***D. teissieri*** |  | ***D. yakuba*** |  | **Hybrids** |  |
|  | Open | Forest | Open | Forest | Open | Forest |
| 200 | 0 | 0 | 141 | 23 | 0 | 0 |
| 650 | 0 | 5 | 241 | 44 | 0 | 0 |
| 1,200 | 14 | 311 | 264 | 30 | 24 | 7 |
| 1,420 | 23 | 201 | 173 | 19 | 44 | 8 |
| 1,650 | 16 | 284 | 145 | 22 | 15 | 2 |
| 1,850 | 18 | 154 | 71 | 6 | 28 | 14 |
| 2,020 | 0 | 50 | 31 | 1 | 2 | 0 |
| 2013 Total | 71 | 1,005 | 1,066 | 145 | 113 | 31 |

We modeled the probability of male *D. teissieri, D. yakuba*, and their hybrids choosing forest habitat over open habitat. We fitted a generalized linear mixed model with binomially distributed error using the ‘*lme4*’ library (Bates et al. 2015). The full model included genotype (either *D. teissieri, D. yakuba*, or their hybrids) and the random effect of block (*N* = 3 blocks). We calculated mean values using parameter estimates from the most likely model, and took the inverse logit of these values to calculate the estimated mean probabilities of choosing forest habitats. To estimate *P*-values, we fitted the same model again using the *mixed* function within the ‘*afex*’ library (Singmann et al. 2015). This approach first fits the full model, and then individual fixed effects are removed to compare the reduced model to the full model. *P*-values were calculated using the likelihood ratio test (LRT) method within *mixed*. These analyses and all others were conducted using the R Statistical Package (version 3.3.1).

### Distributions of *D. yakuba, D. teissieri*, and their hybrids across the ecotone

The male individuals that we recaptured during our mark-recapture experiment enabled us to estimate the rate of hybridization and to assess the distributions of each genotype across the ecotone. Flies collected during this experiment included both dusted flies and any other *D. yakuba* clade flies visiting our traps at the time of recapture. We combined these data with our original capture data to estimate the prevalence of *D. yakuba* genotypes across the ecotone. We first calculated the dilution factor, for each genotype, defined as the number of non-dusted flies captured divided by the number of dusted flies recaptured (White et al. 1982). We then multiplied the number of flies that we originally released by the dilution factor to estimate the census size of each genotype at this single altitudinal sliver of habitat. The estimated census size of hybrids divided by the sum of the three estimated census sizes (*D. yakuba, D. teissieri*, and hybrids) provides an estimate for hybrid prevalence. To assess variation in the prevalence of *D. teissieri* and *D. yakuba* with distance from the center of the ecotone, we fitted logistic models using the *‘MASS’* library.

### Temperature and humidity measurements in the field

We measured the temperature and the humidity of secondary Bioko forest and adjacent open habitat at five of the seven altitudes where flies were sampled (approximately 200, 650, 1,200, 1,650, and 2,020 m). Temperature was measured at 4 times of the day using a Digi Sense field thermometer (Catalog number: 86460-05; Cole-Parmer Instrument Company, Vernon Hills, IL). All measurements were made 10 cm away from the floor. In parallel, and at the same time, we measured air humidity using a field hygrometer (Dwyer MST2-01 Moisture Meter; Michigan City, IN). We recorded these data at five locations per altitude, for each of the two environments, at five altitudes, resulting in a total of 50 measured sites. At each site, we recorded temperature and humidity at 0600, 1200, and 1700 hrs. Each site was measured on three different days for a total of 450 observations.

To assess variation in temperature and humidity, we fitted two linear models, one for temperature and one for humidity. The two models followed the same form and assessed the effect of altitude, type of habitat, and time of the day on each of the two environmental factors. We also included all possible interactions. We compared temperature and humidity at open versus forest habitats using the Tukey’s Honest Significant Difference (HSD) post hoc pairwise comparisons in the *’multcomp’* library (Hothorn et al. 2008).

### Temperature preference in the laboratory

We measured the thermal preferences of *D. teissieri, D. yakuba*, and F_1_(♀*tei* × ♂ *yak*) and F_1_(♀*yak* × ♂ *tei*) hybrids in the laboratory. Flies were randomly placed in a thermocline that consisted of a plexiglass chamber (12 cm wide × 45 cm long × 1 cm high) with an aluminum floor. The thermocline was placed in an 18°C room and heat plates (120 VAC Thermo Scientific Cimarec Hot Plate, Thermo Scientific Cimarec, # UX-04600-01, Waltham, MA, USA) were used to generate a thermal gradient ranging from 18°C to 30°C, with a change in temperature of approximately 2°C every 6 cm. This range of temperatures encompasses the majority of the temperatures flies might regularly experience on Bioko (see below). Flies were allowed to move freely along this gradient over a period of one hour. At the end of each trial we isolated flies into seven chambers, each 10.5 × 6 × 1 cm, by pushing a rod connected to six plexiglass partitions across the width of the chamber. We then recorded the temperature within each partition using a Digi-Sense thermometer equipped with a type-T thermocouple (Cole-Parmer Instrument Co., Chicago, IL; catalog number: 86460-05) and counted the number of flies within each partition. Males and females of each genotype were evaluated separately to avoid sexual attraction that might influence results (*N* = 3 separate replicates per sex).

Because temperature preference was not normally distributed, and remained non-Gaussian after both log (Shapiro Wilk test, *W* = 0.914, *P* < 0.0001) and square root (Shapiro Wilk test, *W* = 0.919, *P* < 0.0001) transformations, we used the *’ARTool’* library to complete analyses of variance on aligned rank transformed data (Kay and Wobbrock 2016). We specifically assessed the effects of genotype, sex, and their interactions on temperature preference. We then used *lsmeans* to conduct post hoc pairwise comparisons (Lenth 2016a).

### Humidity preference in the laboratory

Relative humidity (RH) preferences of *D. teissieri, D. yakuba*, and F_1_(♀*tei* × ♂*yak*) and F_1_(♀*yak* × ♂*tei*) hybrids were evaluated by giving flies the choice of orienting themselves along a humidity gradient. Rows of a 48-well polystyrene tissue culture plate (8 rows by 6 columns; Corning Incorporated, Life Sciences, Tewksbury, MA, USA) were filled with one of three super-saturated salt solutions: LiCl, NaCl, or KH_2_PO_4_. Each of these solutions generates a RH of ~20%, ~70%, and ~85%, respectively, in the headspace above the rows. We filled the wells of two adjacent end rows with LiCl, the three following rows with NaCl, and the three remaining rows with KH_2_PO_4_. This generates a gradient of RH ranging from ~20% to 85%. The top of each plate was covered with 300 micron nylon netting (MegaView Science Co., Ltd. Taichung, Taiwan) and covered with the culture plate lid on top of the mesh; this left ~1 cm for flies to move freely around the plate. This experimental design is a modified approach from Enjin and colleagues (2016). We lightly anesthetized approximately 50, 4 to 7-day old virgin, males or females of a given genotype and placed them along the long axis of a plate. We ran eight plates simultaneously: one plate for each of the parental and hybrid genotypes, with sexes evaluated separately. Flies were allowed to orient themselves for the first hour after which pictures were taken every 15 min, for an additional two hours. This procedure was repeated on four separate days with the position of each plate and the orientation of the ‘low’ and ‘high’ RH end of the plates relative to the room randomized each day. (This avoids confounding effects of non-uniform lighting and other conditions among days in the laboratory.) In total, we assayed ~200 individuals of each genotype and sex. To score preference, we counted the number of flies over each well of the plates, for each of the eight images generated over the 2 hr assay, and summed counts of flies oriented over wells containing the same super-saturated salt solution. All scoring was double blinded.

Because there was no effect of sex on the number of flies choosing a given RH (LRT: χ^2^ = 0.25; *P* = 0.62), sexes were analyzed together. We first tested whether humidity preference varied across genotypes by modeling the mean number of individuals choosing a given RH (Poisson) as a function of genotype, RH, and the interaction between genotype and RH. A dummy ‘plate identity’ variable was included as a random effect to account for among-plate variation such as the location of the plate in the room, slight differences in solution volume within individual wells, or variation in temperature among days. We tested whether each fixed effect influenced the number of flies choosing a given RH using LRTs that compared models that included and excluded each term. We dropped each fixed effect independently, starting with the interaction term. To explicitly test whether different genotypes displayed preferences for different RH, we also modeled the mean number of individuals choosing a given humidity, for each genotype separately, as a function of RH (fixed effect) and plate identity (random effect). We tested for variation in humidity preference using LRTs comparing these genotype-specific models to those lacking the fixed effect of RH. For genotypes that had significant variation in humidity preference we carried out pairwise contrasts between each RH using the *lsmeans* and *pairs* functions of the ‘*lsmeans*’ library (Lenth 2016b).

### Desiccation tolerance in the laboratory

Desiccation resistance was measured by placing ten, 4-day old virgin, females or males in 30 mL empty vials (*N* = 11 vials per sex), which in turn were placed in a glass desiccator with 200 g of Drierite (Sigma Aldrich Catalog number: 7778-18-9; St. Louis, MO) and kept at 21°C (Matute and Harris 2013). The relative humidity was kept under 20% and was measured with a hygrometer. Flies were checked every 30 minutes and the time of death was recorded for each fly.

Differences in desiccation resistance between genotypes were analyzed using a survival analysis and a Cox regression (’rms’ library, Harrell Jr. 2013) using the *cph* function. Plots were generated with the *’survplot*’ library. To assess any effects of body size on desiccation tolerance, we measured the thorax length of 4-day old males and females (*N* = 50 of each sex from each genotype) reared at 24°C. We then fitted a linear model to evaluate the fixed effects of sex and genotype, and the interaction between these factors, using the function *lm* function in the ‘*stats*’ package.

## RESULTS

### Morphological characteristics differ among *D. yakuba, D. teissieri*, and F_1_ hybrids

Before sampling the island of Bioko, we first confirmed whether *D. yakuba*, *D. teissieri*, and F_1_ hybrids could be reliably distinguished based on morphology in the laboratory. We found that while *D. teissieri* males always have at least three strong chitinized anal spines (5.68 ± 1.03 SD), *D. yakuba* males have none, and hybrids have an intermediate number [F_1_(♀*tei* × ♂*yak*), 3.052 ± 0.901 SD; F_1_(♀*yak* × ♂*tei*), 3.196 ± 0.876 SD; *F_3,1996_* = 3356.9, *P* < 0.0001]. All genotypes differed significantly in post hoc comparisons (*P* < 0.021 for all statistically significant comparisons). The lengths of tibia are also intermediate in hybrids [F_1_(♀*tei* × ♂ *yak*), 0.46 ± 0.021 SD; F_1_(♀ *yak* × ♂ *tei*), 0.457 ± 0.02 SD] relative to *D. teissieri* (0.452 ± 0.05 SD) and *D. yakuba* (0.478 ± 0.049 SD) (*F*_3,1996_ = 36.403, *P* < 0.0001)—all groups differed significantly in Tukey’s HSD post hoc pairwise comparisons (*P* <0.037 for all statistically significant comparisons), with the exception of reciprocal hybrids (*P* = 0.203) and the comparison between F_1_(♀ yak × ♂ *tei*) hybrids and *D. teissieri* (*P* = 0.888). In contrast, while the number of teeth in sex combs differed among all groups (*F*_3,1996_ = 220.11, *P* < 0.0001), F_1_(♀ *yak* × ♂ *tei*) hybrids had fewer teeth in their sex combs (4.810 ± 1.328 SD) than did *D. teissieri* (5.238 ± 0.826 SD), *D. yakuba* (6.224 ± 0.955 SD), and F1(♀*tei* × ♂ *yak*) hybrids (6.114 ± 0.952 SD). The latter two genotypes were statistically indistinguishable (*P* = 0.332), but all other post hoc comparisons were significant (*P* < 0.0003 for all statistically significant comparisons). Thus, sex chromosomes and/or the cytoplasmic factors likely influence the number of teeth in sex combs.

We next assessed our ability to reliably identify pure species and hybrids using Mahalanobis distances. Figure 1 shows the distributions for the number of anal spines (Figure 1A), the number of teeth in sex combs (Figure 1B), and the lengths of tibia (Figure 1C) for the four male genotypes. We were able to reliably identify pure species (*D. yakuba:* 100/100, *D. teissieri:* 100/100) and F_1_ hybrids (single class, 196/200). We attempted to gain further resolution and assessed our ability to discriminate between F_1_ (♀*tei* × ♂*yak*) and F_1_(♀*yak* × ♂*tei*) reciprocal hybrids, but due to their similar morphology our assignments were incorrect 53% of the time. Thus, for all later field experiments we treated hybrids as a single class, and relied on subsequent genome sequencing to genotype hybrids.

**FIGURE 1.**
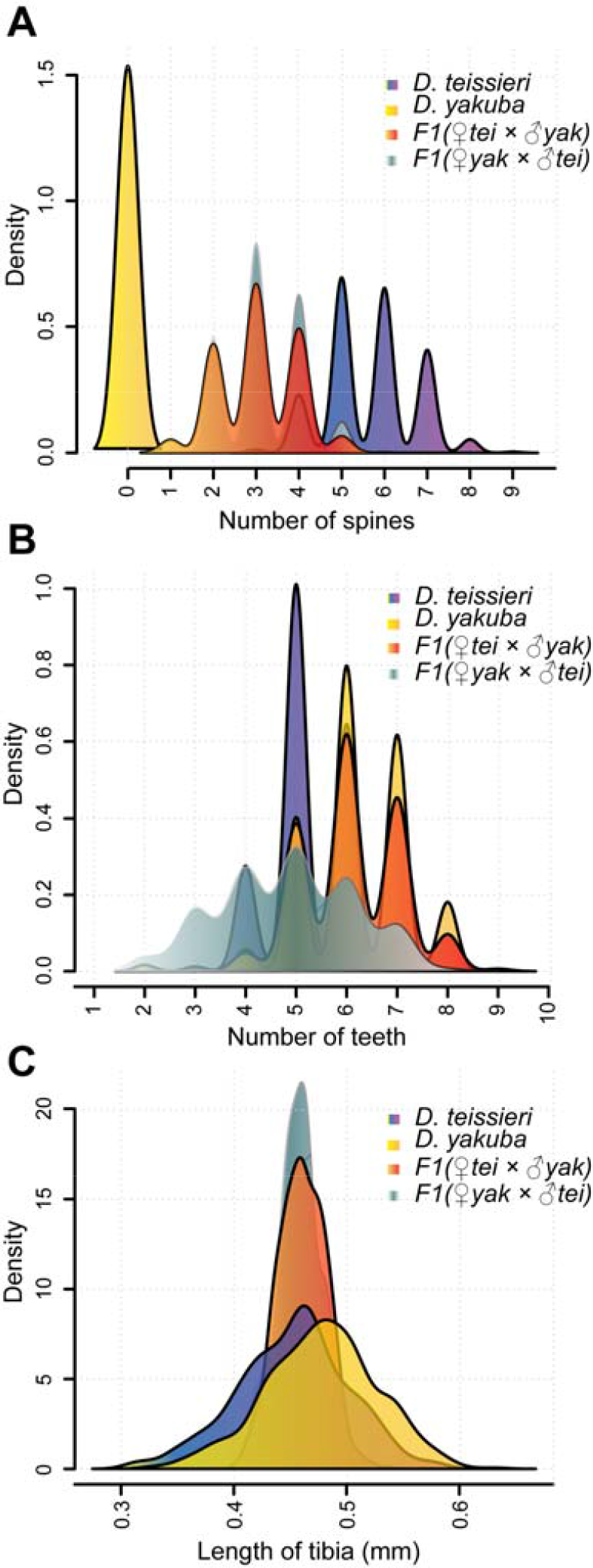
A combination of three phenotypic traits effectively discriminates between *D. yakuba, D. teissieri*, F_1_(♀*tei* × ♂*yak*), and F_1_(♀*yak* × ♂*tei*) genotypes: A) the number of anal spines, B) the number of teeth in sex combs, and C) tibial length (mm). See the text for all statistical analyses. Note that density plots are presented versus histograms for visual clarity, and weights shown for intermediate values are uninformative.

### *Drosophila teissieri* and *D. yakuba* hybridize near the center of the forest-open habitat ecotone

We sampled forest and open habitats across a range of altitudes to determine the geographic distributions of *D. yakuba* and *D. teissieri*, and to identify putative hybrids, in both 2009 and 2013. The results of this sampling are reported in Table 1. Both altitude and habitat type are important factors in separating *D. yakuba* and *D. teissieri* on Bioko. *Drosophila teissieri* was found mostly in higher altitude forests, while *D. yakuba* was found mostly in open areas at all altitudes (200 - 2,020 m), although *D. yakuba* was most common at lower altitudes where the majority of human settlements are located (Table 1). This strong association between habitat type and species presence suggests that *D. teissieri* and *D. yakuba* are largely geographically separated on Bioko. Nevertheless, based on Mahalanobis distances we identified hybrids at the boundary of forest and open habitat, but only at altitudes where *D. teissieri* and *D. yakuba* co-occur (Table 1).

### Hybrids on Bioko are F_1_ progeny of *D. yakuba* females and *D. teissieri* males

To confirm our field genotype assignments based on Mahalanobis distances alone, we sequenced a subset of the male individuals sampled from Bioko, including putative hybrids. All 19 sequenced hybrid males are heterozygous for *D. yakuba* and *D. teissieri* ancestry across their autosomes and hemizygous for the *D. yakuba X* (Figure 2). This indicates that these hybrids are the sons of *D. yakuba* females and *D. teissieri* males. There are many small regions for which genotypes could not be confidently called (gray blocks in Figure 2). Taken together, these data confirm our discovery of hybrids on Bioko based on Mahalanobis distances in the field, but these data also suggest that F_1_(♀*tei*× ♂ *yak*) and advanced-generation hybrids may not occur on Bioko.

**Figure 2:**
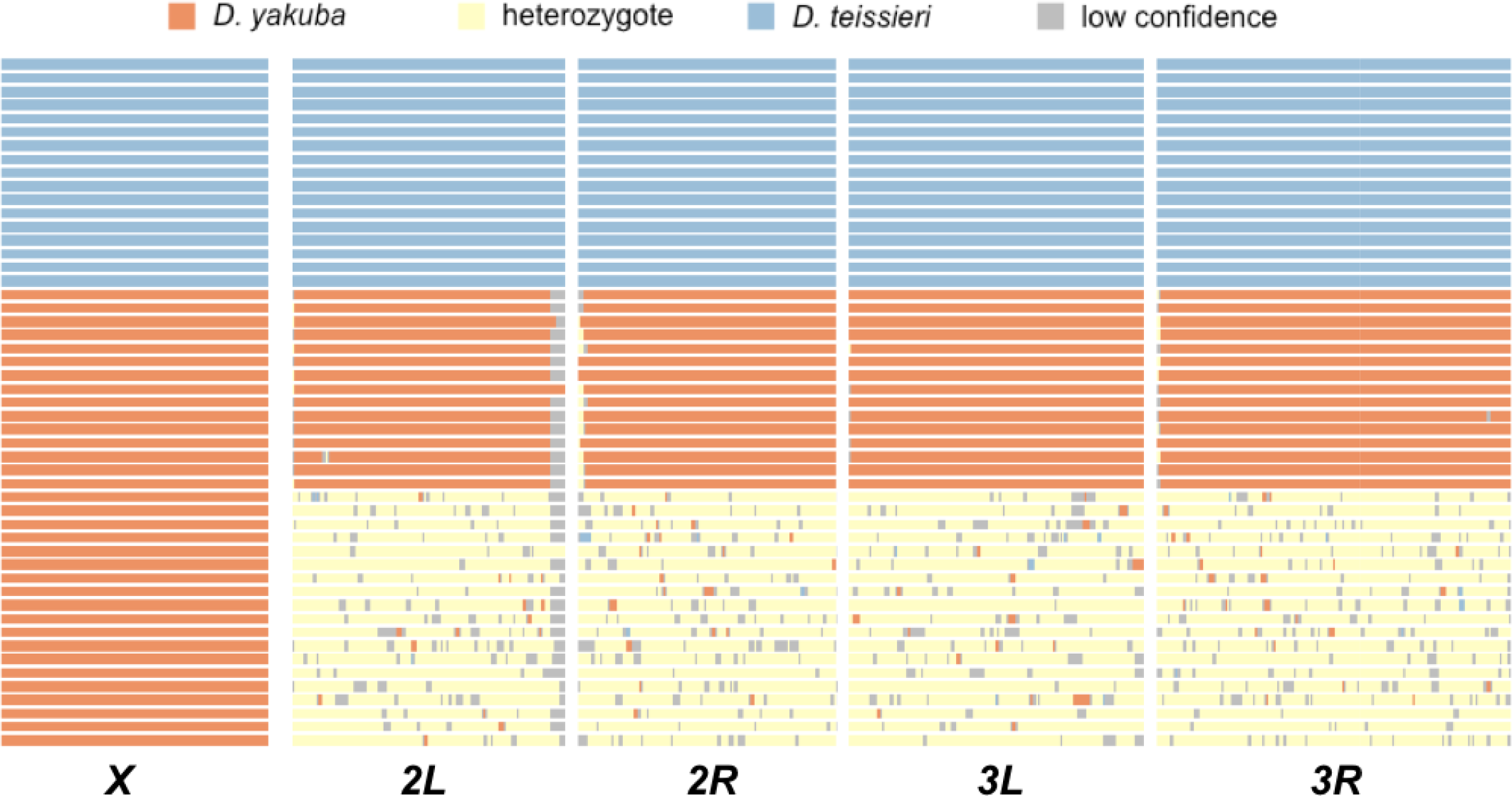
Genomic sequencing confirms that F_1_(♀*yak* × ♂*tei*) hybrids occur on Bioko. Chromosome view of inferred ancestries of wild-caught male *D. yakuba* (orange), *D. teissieri* (blue), and predicted hybrids based on Mahalanobis distances (yellow) are plotted. Chromosomes are denoted at the bottom of the plot. All individuals classified as hybrids in the field were heterozygous across the autosomes for both parental genotypes. Ancestry on the *X* chromosome reflects hemizygous genotypes and indicates that hybrids are F_1_ progeny of *D. yakuba* females and *D. teissieri* males. There are many small regions for which genotypes could not be confidently called (gray).

### *Drosophila teissieri* prefers forests, but *D. yakuba* and hybrid genotypes prefer open habitats

We next characterized the habitat preferences (forest vs. open habitat) of *D. yakuba, D. teissieri*, and their hybrids by dusting flies with UV powder and releasing flies from the center of the forest-open habitat ecotone at 1,650 m. This experiment included the 1,076 *D. teissieri*, 1,211 *D. yakuba*, and 144 hybrid males that we sampled across the altitudinal transect in 2013 (Table 1). In total, we recaptured 91 (8.45%) *D. teissieri* males, 137 (11.31%) *D. yakuba* males, and 43 (30%) hybrid males that were released. In all cases, genotypes assigned to the pure species class had motile sperm and those assigned to the hybrid class did not, as we predicted. Recapture experiments revealed that the estimated mean probability of choosing forest habitat differed among *D. teissieri* (0.97), *D. yakuba* (0.08), and hybrids (0.32) (LRT: *P* < 0.0001). Approximately 3% of *D. teissieri*, 15% of *D. yakuba*, and 19% of hybrids were recaptured at the center of the forest-open habitat ecotone (Figure 3). Together these results indicate that genotypes differ in their habitat preference, but hybrid preference [predicted to be F_1_(♀*yak* × ♂*tei*) genotypes] for open areas more closely aligns with that of *D. yakuba*. This suggests *D. yakuba* alleles involved in habitat choice are dominant or semi-dominant to *D. teissieri* alleles.

**FIGURE 3.**
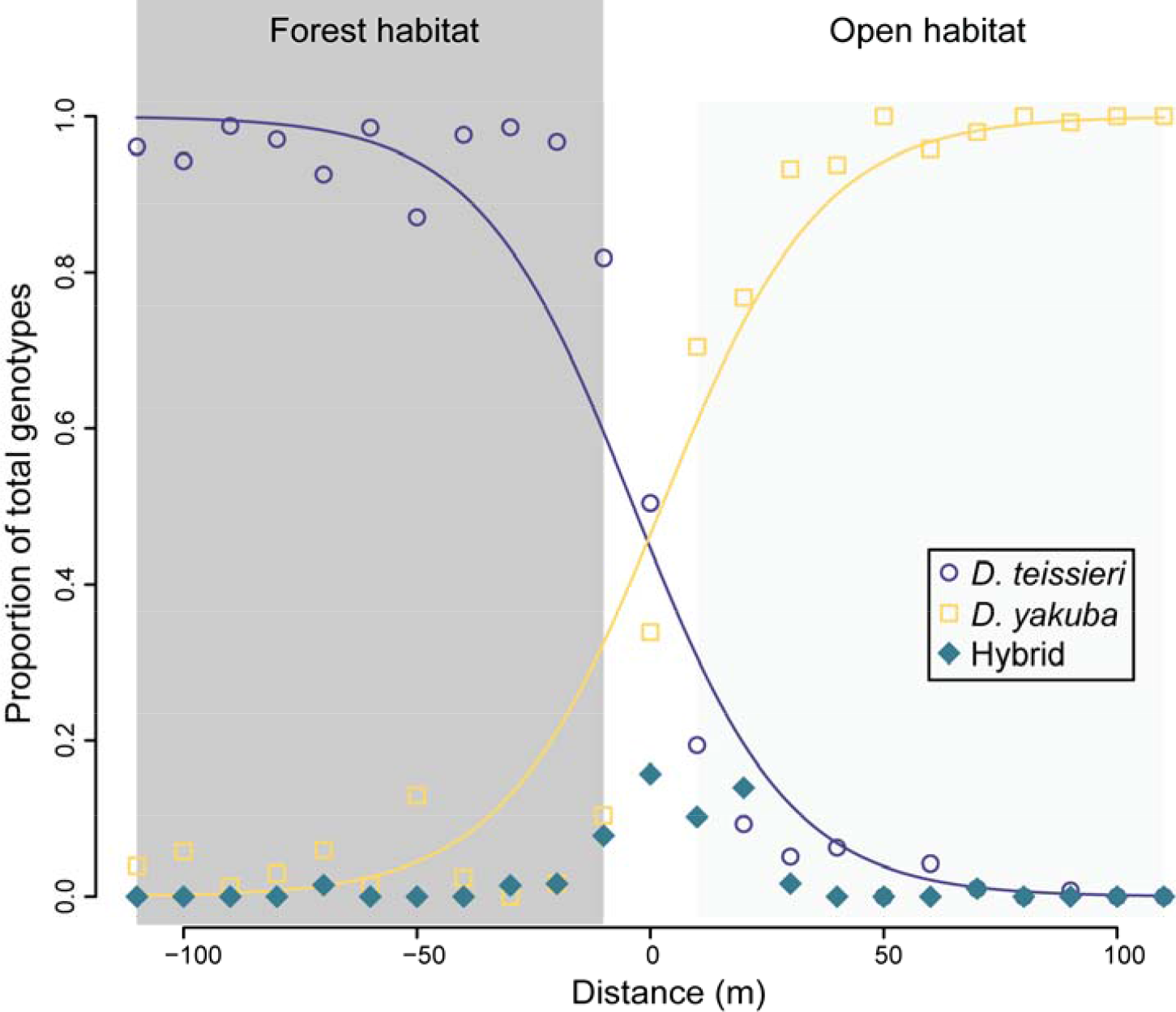
*Drosophila teissieri* occurs in forests, *D. yakuba* occurs in open areas, and hybrids occur in a very narrow region centered on the forest-open habitat ecotone. Plotted is the proportion of each genotype sampled from each trap during the recapture portion of our mark-recapture experiment. This includes a total of 951 *D. teissieri* males, 1,172 *D. yakuba* males, and 59 hybrid males. These sample sizes include the 91 *D. teissieri* males, 137 *D. yakuba* males, and 43 hybrid males that were dusted and recaptured, in addition to any other *D. yakuba* clade individuals visiting the traps. More than 98% of all hybrids were captured within 30 m of the release point at the center of the ecotone (0 m), suggesting that the hybrid zone is very narrow. Fitted lines from logistic regressions are plotted for *D. yakuba* and *D. teissieri*. The prevalence of each varies with distance from the center of ecotone (P < 0.0001) with *D. teissieri* mostly found in forest habitat and *D. yakuba* mostly found in open habitat.

### Hybridization is relatively rare and the hybrid zone is narrow

The recapture experiments enabled us to estimate the commonness of *D. yakuba, D. teissieri*, and their hybrids across the ecotone. In total, we captured 951 *D. teissieri*, 1,172 *D. yakuba*, and 59 hybrid males during the recapture portion of our mark-recapture experiment. This included 91 *D. teissieri*, 137 *D. yakuba*, and 43 hybrid (dusted) males that we released. We estimate that with our measured recapture rates (11.31% for *D. yakuba* and 8.45% for *D. teissieri*), the number of flies at this 1,650 m site is close to 1 × 10^4^ for each species. The dilution factor (non-dusted flies captured/dusted flies recaptured) was 7.55 for *D. yakuba* and 9.45 for *D. teissieri*. Thus, the estimated census sizes of *D. yakuba* (*N* ~ 9,150) and *D. teissieri* (*N*~ 10,170 for *D. teissieri*) are about 10 times larger than the total number of flies we initially caught (*N* = 1,076 *D. teissieri* and *N* = 1,211 for *D. yakuba*). These estimates only constitute the approximate number of flies at a single altitudinal sliver of habitat.

Hybrids seem to be relatively rare and occur in a narrow zone centered on the forest-open habitat ecotone. In total, we recaptured 43 dusted hybrids plus an additional 16 hybrids visiting the traps during the recapture portion of our mark-recapture experiment. (73% of the flies caught on the second attempt had already been captured.) Thus, we estimate there are about ~55 hybrids at our sampled location, and that hybrids constitute about 0.3% of the total number of *D. yakuba* clade individuals in this area. These are clearly rough estimates, but they suggest that hybrids are not common. 98% of the hybrids that we recaptured traveled less than 30 m from the ecotone, suggesting that this hybrid zone is very narrow (Figure 2). *Drosophila teissieri* was mostly absent from the open habitat and its prevalence increased with distance into the forest habitat (*z*-value = -22.147, df = 21, *P* < 0.0001). In contrast, *D. yakuba* was mostly absent from forest habitat and its prevalence increased (*z*-value = 22.267, df = 21, *P* < 0.0001) with distance into the open habitat (Figure 2). Taken together these data indicate a sharp transition in the distributions of *D. teissieri* and *D. yakuba* at the center of the forest-open habitat ecotone where they come into contact and produce F_1_(♀*yak* × ♂*tei*) hybrids; these hybrids were never sampled more than 70 m from the center of the ecotone.

### Abiotic conditions differ among altitudes and between habitats on Bioko

*Drosophila yakuba, D. teissieri*, and their hybrids occur and co-occur at only certain altitudes, and differ in the preferences for forest and open habitat. Because temperature and humidity influence the physiology and geographical distributions of *Drosophila* species (Hoffmann and Kellermann 2006; Hoffmann and Weeks 2007; Kellermann et al. 2009; Kellermann et al. 2012a; Kellermann et al. 2012b; Cooper et al. 2014; Adrion et al. 2015), we measured these factors in forest and open habitats near the center of the ecotone, at five altitudes (approximately 200, 650, 1,200, 1,650, and 2,020 m), and at three times of the day (0600, 1200, and 1700 hrs). In general, and as expected, both temperature and humidity decreased with altitude (Table 2, Figure 4). Forest sites were more humid than were open habitats, and forest sites had lower temperatures at all times of day, with the exception of 0600 at low altitudes Figure 4A. Means, standard deviations, and statistics are presented in Table 3. These data suggest that on average *D. teissieri* and *D. yakuba* experience different temperature and humidity conditions in the field, and that hybrids most often experience relatively warm and dry conditions in their preferred open habitat.

**FIGURE 4.**
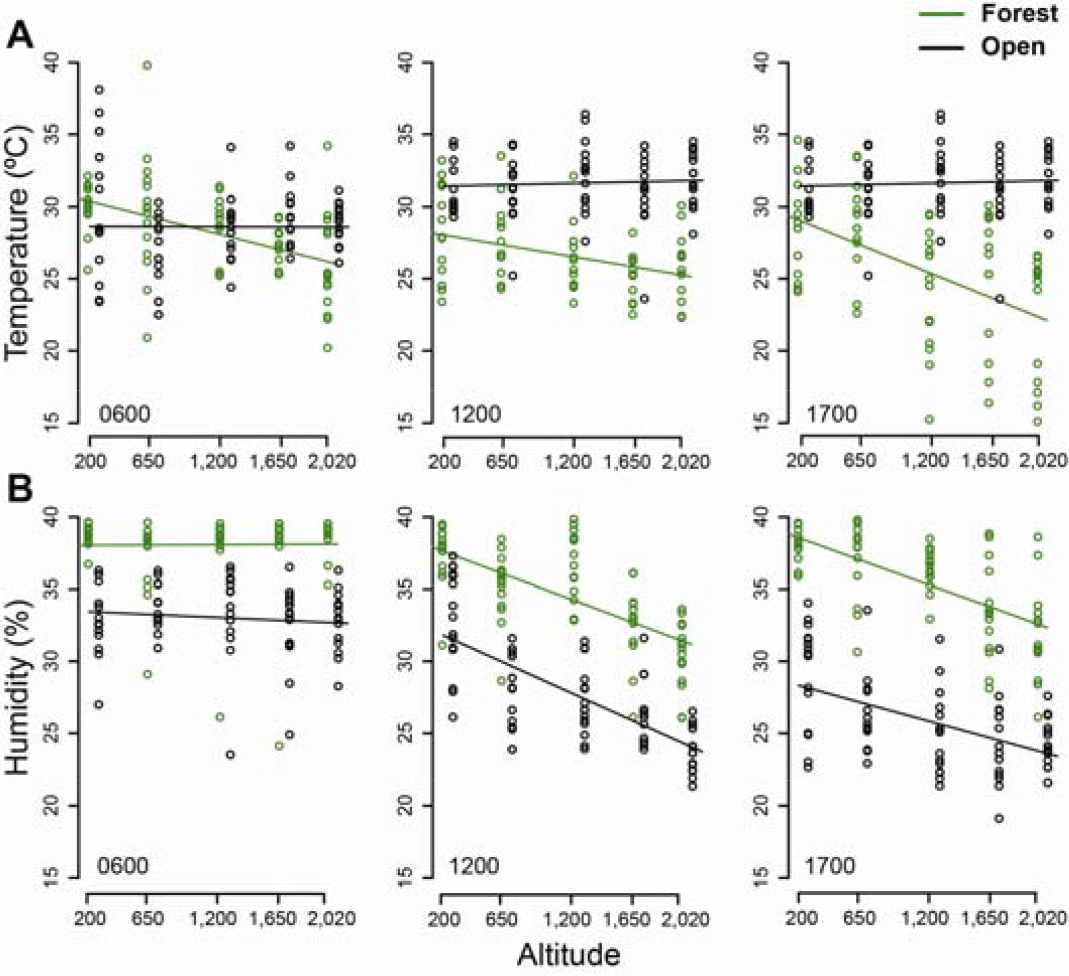
Abiotic conditions at different altitudes vary between forests and open habitats, and at different altitudes on Bioko. Each panel reports environmental measurements for a specific time of the day labeled in the bottom left corner of each panel. (A) With the exception of low altitudes in the early morning, temperature (°C) is generally higher in open areas than in forests. (B) In contrast, humidity (%) is generally higher in forests than in open habitats. Regression lines for effects of altitude on each environmental condition, at each time of day, are plotted. Values and lines for open habitats are offset for visual clarity. The distributions of *D. yakuba* and *D. teissieri* and their hybrids, are reported in Table 1. Statistical analyses are presented in Table 2.

**TABLE 2.** ANCOVA reveals significant effects of both altitude and habitat (open vs. forest) on both temperature and humidity across the island of Bioko.

| **Environmental condition** | **Factor** | **df** | ***F*-value** | ***P*-value** |
| --- | --- | --- | --- | --- |
| Temperature | Altitude | 1 | 19.19 | < 0.0001 |
|  | Habitat | 1 | 177.79 | < 0.0001 |
|  | Altitude × Habitat | 1 | 24.45 | < 0.0001 |
| Humidity | Altitude | 1 | 519.55 | < 0.0001 |
|  | Habitat | 1 | 171.54 | < 0.0001 |
|  | Altitude × Habitat | 1 | 35.56 | < 0.0001 |

**TABLE 3.** Mean and standard deviation of temperature and humidity in the island of Bioko. For these calculations measurements at different altitudes and different times of the day were pooled. *t*-values and *P*-values of Tukey’s HSD pairwise comparisons between forest and open habitats are reported for both temperature and humidity.

|  | **Temperature** |  | **Humidity** |  |
| --- | --- | --- | --- | --- |
|  | **Mean (°C)** | **SD (°C)** | **Mean (%)** | **SD (%)** |
| Forest habitat | 26.70 | 3.58 | 83.95 | 14.09 |
| Open habitat | 30.61 | 2.89 | 55.24 | 17.94 |
| Tukey's HSD | \|t-value\|= 2.357; *P* = 0.019 |  | \|t-value\|= 2.634; *P* = 0.009 |  |

### Hybrids have a maladaptive combination of parental phenotypes

The F_1_(♀*yak* × ♂*tei*) hybrids we sampled on Bioko behaviorally prefer warm and dry open areas 68% of the time in the field—these areas are also chosen by *D. yakuba* 92% of the time. In contrast, *D. teissieri* chooses these areas only 3% of the time. We next sought to determine whether these preferences exist in the laboratory, and if so, whether the physiological tolerances of *D. yakuba, D. teissieri*, and hybrids to low humidity conditions corresponds to their behavioral preferences in the field.

***Drosophila yakuba* and F_1_ hybrids prefer warmer temperatures than *D. teissieri***. We assessed the temperature preference of *D. yakuba, D. teissieri*, and their F_1_ hybrids in the laboratory. Temperature preferences for the eight lab-produced genotypes [(2 pure species + 2 reciprocal hybrids) × 2 sexes] are shown in Figure 5. The mean temperature preference of *D. teissieri* (21.285°C ± 2.905 SD, *N* = 1,200) was approximately 13% lower than *D. yakuba* (24.51°C ± 3.68 SD, *N* = 1,200), 12% lower than F_1_(♀*tei* × ♂*yak*) (24.12°C ± 3.72 SD, *N*= 1,200), and 10% lower than F_1_(♀*yak* × ♂*tei*) hybrids (23.74°C ± 3.82 SD, *N* = 1,200) (*F_3,4792_* = 209.927, *P* < 0.0001). Tukey’s HSD post hoc comparisons revealed the temperature preference of F_1_(♀*tei* × ♂*yak*) hybrids differed from *D. teissieri* (*P*< 0.0001), but not *D. yakuba* (*P* = 0.05). In contrast, F_1_(♀*yak* × ♂*tei*) hybrids differed from both *D. teissieri* (*P* < 0.0001) and *D. yakuba* (*P* < 0.0001). These patterns stem from a slight, but statistically significant, difference between the two reciprocal hybrids (*P* = 0.03). Across all genotypes females preferred higher temperatures (23.87°C ± 3.70 SD) than did males (22.95°C ± 3.78 SD) *(F_1,4792_* = 57.361, *P* < 0.0001). Finally, we found a modest, but statistically significant, interaction between genotype and sex (*F_3,4792_* = 3.207, *P* < 0.022). These results indicate that F_1_ hybrid genotypes have a preference for warm temperatures that is similar to *D. yakuba* in the laboratory, in addition to the field, suggesting that *D. yakuba’s* preference for high temperatures is dominant or semi-dominant to *D. teissieri’s* preference for cool temperatures.

**FIGURE 5.**
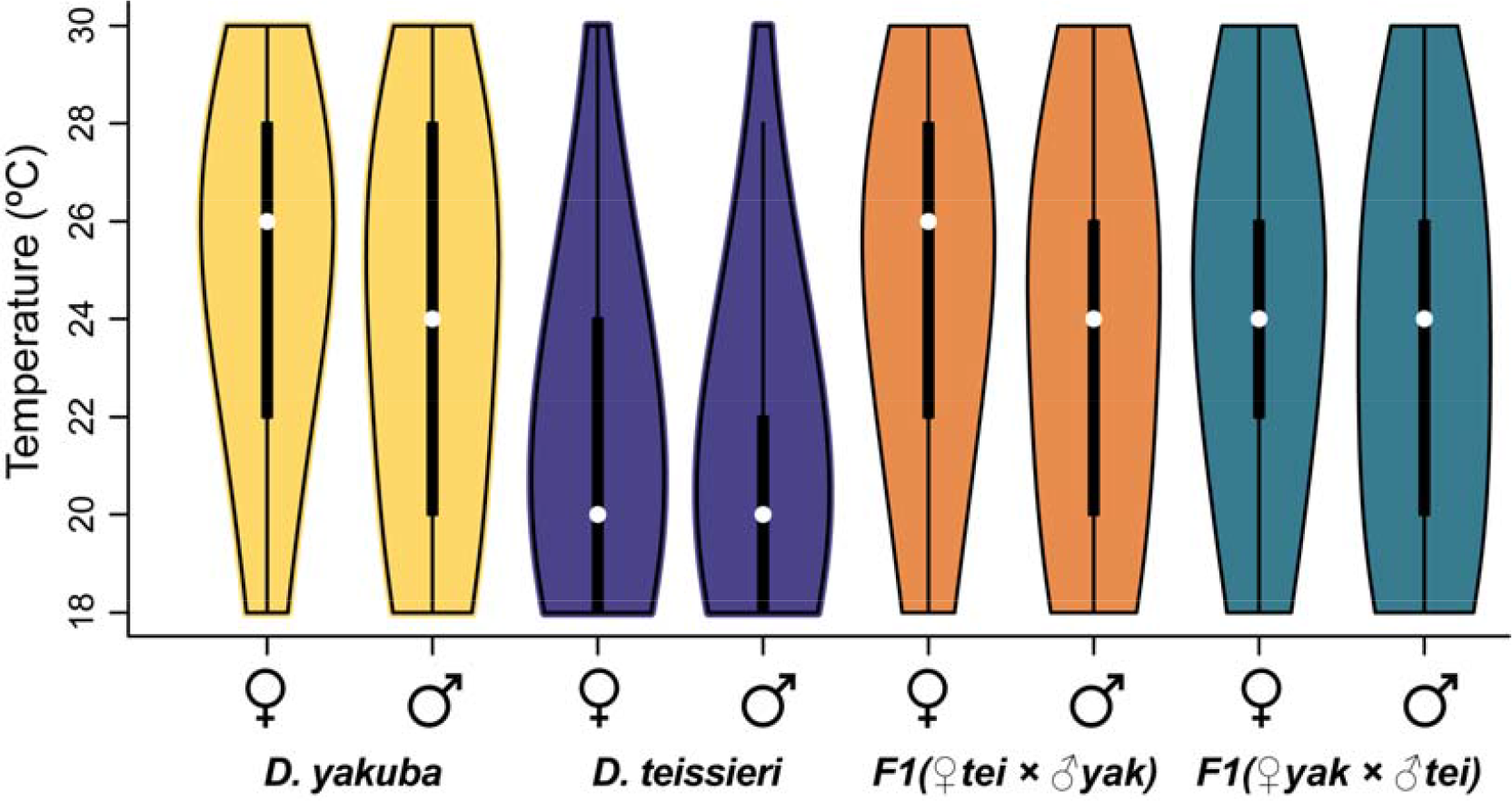
In the laboratory, *D. yakuba*, F_1_(♀*tei* × ♂*yak*), and F_1_(♀*yak* × ♂*tei*) genotypes prefer warmer temperatures than does *D. teissieri*, as measured in a laboratory thermocline. Sexes of each of the four genotypes are presented independently. *Drosophila teissieri* shows a significantly lower temperature preference than the other three genotypes (*F_3,4792_* = 209.927, *P* < 0.0001). See text for pairwise contrasts.

***Drosophila yakuba* and F_1_ hybrids prefer dryer conditions than *D. teissieri***. We next assessed the humidity preference of *D. yakuba, D. teissieri*, and their F_1_ hybrids. We did not observe a significant interaction between genotype and RH (LRT: χ^2^ = 7.27; *P* = 0.30); however, both genotype and RH affected the number of individuals observed at a given location along the gradient (LRTs: χ^*2*^ = 7.27; *P* = 0.30; χ^2^ =7.27; *P* = 0.30, respectively). When analyzed individually, *D. teissieri* did not show a preference for a specific RH (LRT: χ^2^ = 4.1; *P* = 0.13; Figure 6), while *D. yakuba* preferred low RH (20% RH; LRT: χ^2^ = 15.2; *P* < 0.001; Figure 6). Both types of F_1_ hybrids behaved similarly to *D. yakuba* in that they tended to prefer lower humidity [LRTs; F_1_(♀*yak* × ♂♂*tei*): χ^2^ = 7.8; *P* = 0.02; F_1_(♀*tei* × ♂*yak*): χ^2^ = 24.4; *P* < 0.0001; Figure 6], but F_1_(♀*yak* × ♂*tei*) hybrids show a weaker preference than F_1_ (♀*tei* × ♂*yak*) hybrids. This is indicated by the only marginally significant pairwise contrasts of the number of F_1_(♀*yak* × ♂*tei*) individuals at different humidity (*P* = 0.06 for both 20% - 70% RH and 20% - 85% RH), but highly significant contrasts for F_1_(♀*tei* × ♂*yak*) hybrids (*P* = 0.0004 for both 20% - 70% RH and 20% - 85% RH). These data indicate that F_1_ hybrid genotypes have a preference for low humidity that is similar to *D. yakuba*, suggesting that *D. yakuba*’s preference for low humidity is dominant or semi-dominant to *D. teissieri’s* lack of preference.

**FIGURE 6.**
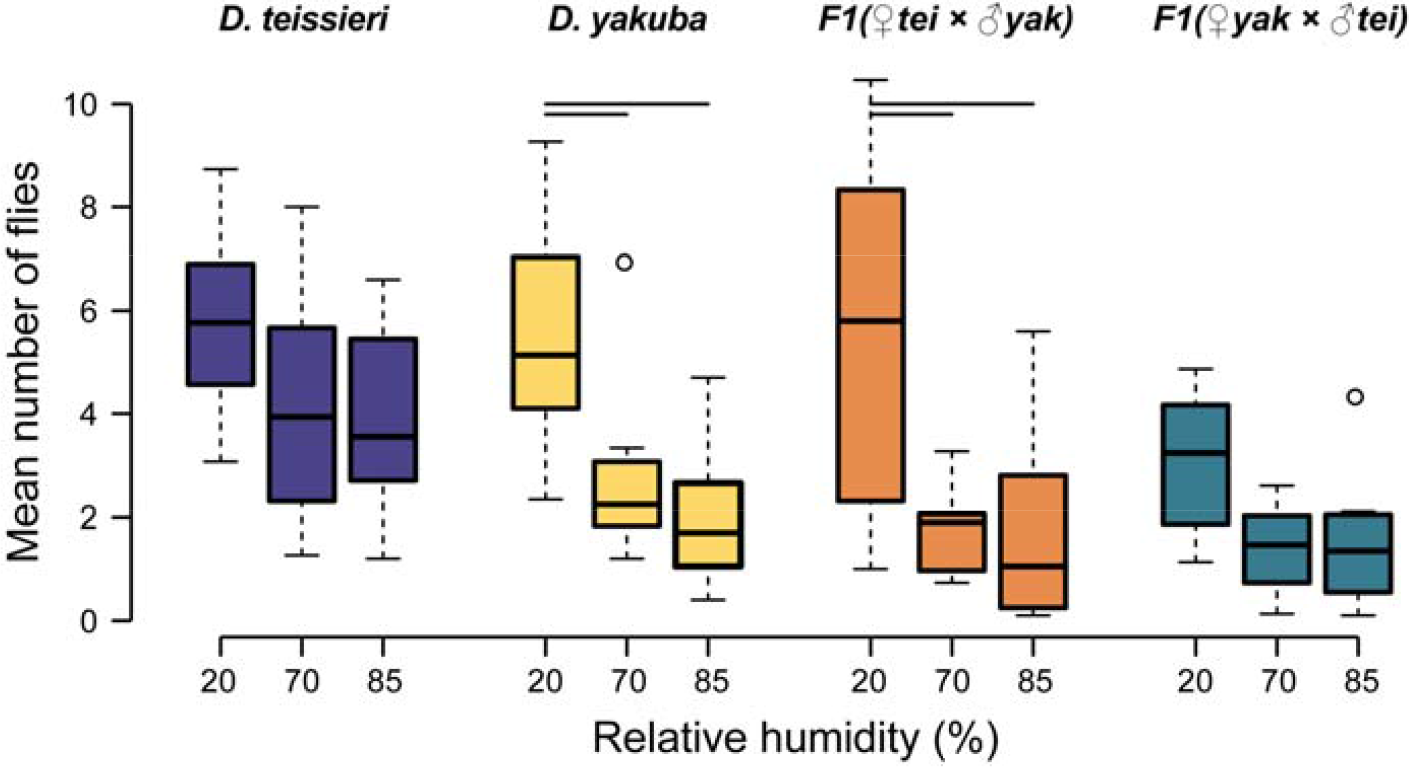
*Drosophila teissieri* prefers higher relative humidity (%) than does *D. yakuba*, F_1_(♀*tei* × ♂*yak*), and F_1_(♀*yak* × ♂*tei*) genotypes as measured in the laboratory. Like *D. yakuba*, both types of F_1_ hybrids tended to prefer lower humidity [LRTs; F_1_(♀*yak* × ♂*tei*): χ^2^ = 7.8; *P* = 0.02; F_1_(♀*tei* × ♂*yak*): χ^2^ = 24.4; *P* < 0.0001]. Horizontal bars represent significant pairwise contrasts (*P* < 0.01) between humidity conditions fit separately for each genotype (see text for details).

**F_1_ hybrids have low desiccation tolerance that closely resembles that of *D. teissieri***. Finally, we assessed whether *D. yakuba, D. teissieri*, and their F_1_ hybrids differ in physiological tolerance to osmotic stress. The body size of all genotypes was similar suggesting that any differences in desiccation tolerance do not depend on size (*F*_3,392_ = 2.4335, *P* = 0.065). As expected, males were more prone to suffer from desiccation than were females (~between 1% and 10% less tolerant depending on the genotype; Cox hazards regression, sex effect: χ^2^ = 8.39, df = 1, *P* = 0.004)(Table S2)—the smaller relative body size of males likely influences this difference (*F*_1,392_ = 295.25, *P* < 0.0001). The largest differences in desiccation tolerance were observed between genotypes with *D. yakuba* having the highest desiccation tolerance (Cox hazards regression, genotype effect: χ^2^ = 143.55, df = 3, *P* < 0.0001) (Figure 7). The tolerance of *D. teissieri* to osmotic stress (4.73 hr ± 1.69 SD) is 40% lower than the tolerance of *D. yakuba* (6.63 hr ± 1.54 SD), consistent with the distribution of these species in nature. Together, hybrids from both reciprocal crosses have approximately 28% lower desiccation tolerance than *D. yakuba*, despite behaviorally preferring such conditions. F_1_(♀*yak* × ♂*tei*) hybrids that occur naturally on Bioko had the lowest desiccation tolerance of all genotypes (4.50 hr ± 1.47 SD), however, these hybrids did not differ statistically from F_1_(♀*tei* × ♂*yak*) hybrids (4.76 hr ± 1.50 SD). This suggests little influence of the cytoplasm or sex chromosomes on desiccation tolerance. Finally, F_1_(♀*yak* × ♂*tei*) hybrids have significantly lower desiccation tolerance than does *D. teissieri*, but F_1_(♀*tei* × ♂*yak*) hybrids do not differ statistically from *D. teissieri*. Means, standard deviations, sample sizes, and statistical comparisons are reported in Table 4. Taken together, these results indicate that *D. teissieri* alleles influencing desiccation tolerance are dominant or semi-dominant to *D. yakuba* alleles. Thus, F_1_ hybrids, and especially F_1_(♀*yak* × ♂*tei*) hybrids, are predicted to be physiologically ill-equipped to cope with their behaviorally preferred conditions.

**FIGURE 7.**
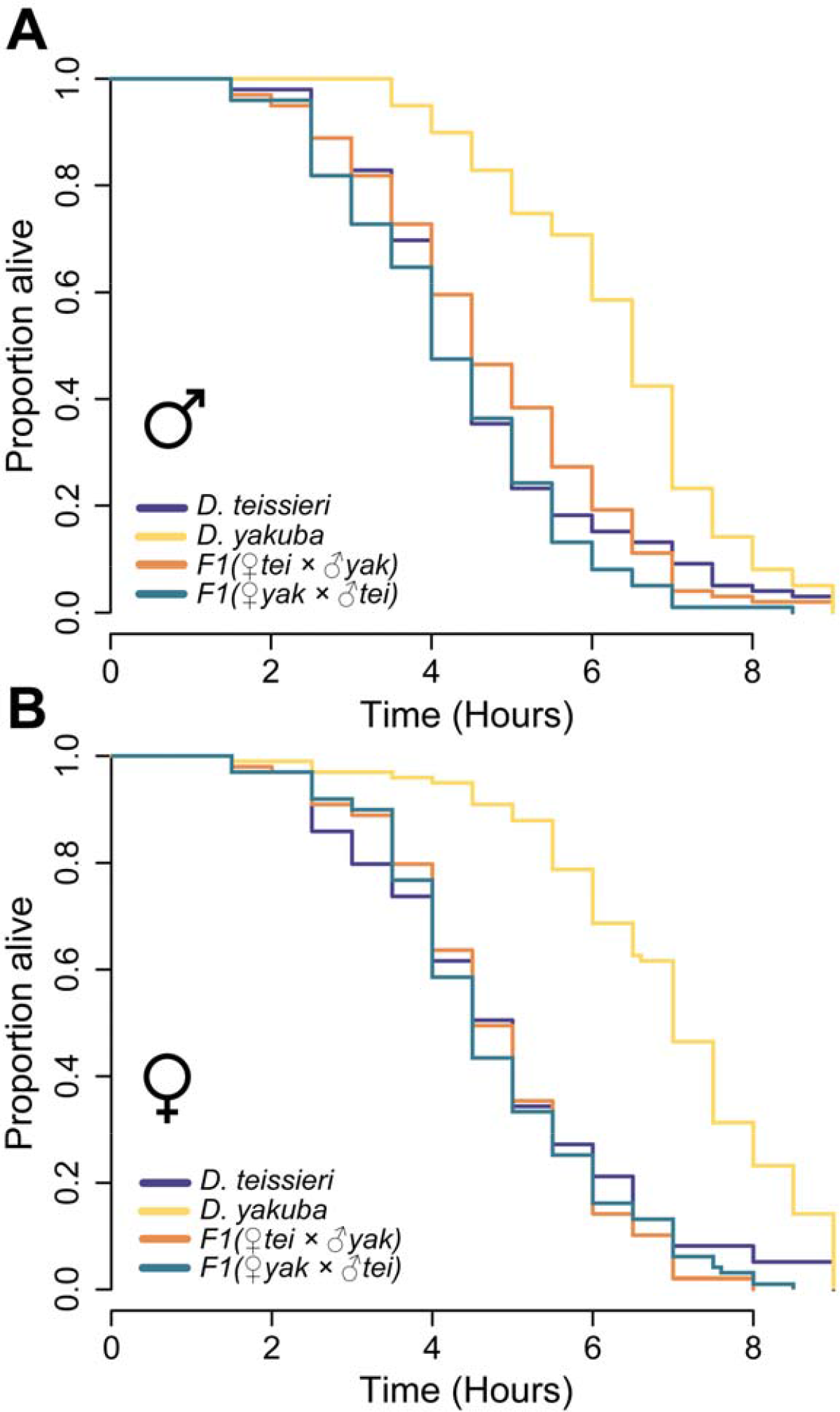
Both male (A) and female (B) *D. yakuba* individuals have higher desiccation tolerance than do *D. teissieri*, F_1_(♀*tei* × ♂*yak*), and F_1_(♀*yak* × ♂*tei*) genotypes. Across sexes F_1_(♀*yak* × ♂*tei*) have the lowest desiccation tolerance indicating that these hybrids are ill-equipped to cope with the dry environments they prefer in the laboratory and in nature. Summary statistics are reported in Table S2 for males and females. Table 4 reports summary statistics and the results of Tukey’s HSD post hoc pairwise comparisons between genotypes.

**TABLE 4.** Desiccation resistance of *D. yakuba, D. teissieri*, and the two reciprocal hybrids—sexes were pooled. Sample sizes (*N*), mean time in hours until death, and standard deviations (SD) are reported. The last four columns show our statistical analyses of particular comparisons as a 4 × 4 matrix. The upper triangular matrix shows the z-values from Tukey’s HSD pairwise comparisons, and the lower triangular matrix shows associated *P*-values. Statistically significant differences (*P* < 0.05) are indicated in boldface.

| **Genotype** | ***N*** | **Mean** | **SD** | *D. yakuba* | *D. teissieri* | $F_1(\text{♀}yak \times \text{♂}tei)$ | $F_1(\text{♀}tei \times \text{♂}yak)$ |
| --- | --- | --- | --- | --- | --- | --- | --- |
| *D. yakuba* | 198 | 6.63 | 1.53 |  | -8.64 | 11.11 | -9.45 |
| *D. teissieri* | 198 | 4.73 | 1.69 | < **0.001** |  | 2.67 | 0.93 |
| $F_1(\text{♀}yak \times \text{♂}tei)$ | 198 | 4.50 | 1.47 | <**0 .001** | < **0.038** | | 1.77 |
| $F_1(\text{♀}tei \times \text{♂}yak)$ | 198 | 4.76 | 1.50 | <**0.001** | 0.789 | 0.29 | |

## DISCUSSION

While challenging, *D. teissieri* and *D. yakuba* can be hybridized under laboratory conditions (Turissini et al. 2015), and notably, both F_1_(♀*yak* × ♂*tei*) and F_1_(♀*tei* × ♂*yak*) females are fertile (hybrid males are sterile) making it possible for female hybrids to backcross with parental species males. Interestingly, similarly diverged pairs of *D. melanogaster* subgroup species produce inviable or sterile hybrids in the laboratory (e.g., *D. melanogaster-D. simulans* hybrids, Ks=0.10; Matute et al. 2010). Whether *D. teissieri* and *D. yakuba* currently hybridize in nature has remained unknown, but here we have presented field and genomic evidence that *D. yakuba* females and *D. teissieri* males hybridize on the island of Bioko. These species are estimated to have diverged ~3.0mya my ago making this the most diverged pair in the genus with a contemporary hybrid zone (Turissini and Matute in revision; Turelli, Conner, Turissini, Matute, and Cooper unpublished). Hybridization occurs above ~1,200 m in the Bioko highlands near the center of the forest-open habitat ecotone where we estimate that hybrids comprise 0.3% of all sampled *D. yakuba* clade flies. Finally, we sampled hybrids in both 2009 and 2013, and 98% of the hybrids we identified were sampled within 30 m of the center of the ecotone, suggesting some stability of this narrow hybrid zone (sensu, Key 1968).

Multiple lines of evidence suggest that the F_1_(♀*yak* × ♂*tei*) hybrids that occur on Bioko have a maladaptive combination of parental traits. In the laboratory, *D. teissieri* prefers relatively cool and high humidity conditions, while *D. yakuba* prefers warm and low humidity conditions. Each species has physiology that matches their behavioral preference: *D. yakuba* has high and *D. teissieri* has low desiccation tolerance. In contrast, hybrids prefer warm and dry conditions, but they have low desiccation tolerance. In the field, the estimated mean probability of hybrids choosing the relatively warm and dry open habitat also preferred by *D. yakuba* is 0.68. This indicates that in hybrids *D. teissieri* allele(s) underlying low desiccation tolerance are dominant or semi-dominant to *D. yakuba* alleles, but the *D. yakuba* allele(s) underlying behavioral preference for warm and dry conditions are dominant or semi-dominant to *D. teissieri* alleles. We predict that this results in a maladaptive combination of parental phenotypes in hybrids.

There are other fitness costs that likely contribute to the maintenance of this narrow hybrid zone. First, all hybrid males are sterile (Turissini et al. 2015). Second, *D. yakuba-D. teissieri* hybrids have behavioral defects that compromise their foraging behavior in ways predicted to reduce hybrid fitness (Turissini et al. 2017). Third, there is some hybrid inviability exacerbated by *Wolbachia* infections that cause cytoplasmic incompatibility (CI) in interspecific crosses between *Wolbachia*-uninfected (U) females and *Wolbachia*-infected (I) males. Females infected by *Wolbachia* are protected from this interspecific CI and the associated reductions in egg hatch. However, the frequencies of *Wolbachia* infections are temporally heterogeneous on São Tomé for both *D. santomea* and *D. yakuba*, and heterogeneous between São Tomé and Bioko populations of *D. yakuba*, suggesting that the proportion of females susceptible to CI fluctuates. While this could contribute to postzygotic RI between *D. yakuba* clade species, we find it unlikely such contributions are substantial (see Cooper et al. 2017 for a discussion). Importantly, the strength of interspecific CI is similar for (U♀*tei* × I♂*yak*) and (U♀*yak* × I♂*tei*) crosses (Table 8, Cooper et al. 2017), indicating that *Wolbachia* infections likely do not explain the paucity of F_1_(♀*tei* × ♂*yak*) hybrids on Bioko. The barriers that limit the production of F_1_(♀*tei* × ♂*yak*) hybrids in nature remain unknown, but laboratory experiments have not revealed any obvious differences in the fertilization success or sterility/viability between F_1_ hybrids produced by reciprocal crosses (Turissini et al. 2015; Cooper et al. 2017), as observed in other systems (i.e., “Darwin’s corollary”, Turelli and Moyle 2007).

A maladaptive combination of traits in F_1_ hybrids provides a possible mechanism for both the lack of advanced-generation hybrids on Bioko and limited introgression in the *D. yakuba* clade. Both mt and *Wolbachia* genomes show divergence among all three *D. yakuba* clade species (Turelli, Conner, Turissini, Matute, and Cooper unpublished), indicating that cytoplasmic introgression is less common than originally described (Monnerot et al. 1990; Llopart et al. 2005b; Bachtrog et al. 2006). This discrepancy seems to be due to the number of loci included in different analyses and on the methods used to assess amounts of introgression. For example, the most recent published whole genome analysis of *D. yakuba* and *D. santomea* mtDNA suggested pervasive introgression (Llopart et al. 2014), but this study relied on IMa2 (Hey and Nielsen 2004, 2007), which is known to provide false positives for migration (Cruickshank and Hahn 2014; Hey et al. 2015).

Only two other hybrid zones have been identified in the *Drosophila*, and both of these examples are for recently diverged sister species in the *D. melanogaster* subgroup. The sister species *D. simulans* and *D. sechellia* hybridize in the central Seychelles. F_1_ hybrid males between these two species have been found in ripe *Morinda* ripe fruits. Such evidence indicates that the ability to breed on *Morinda* could be transferred from *D. sechellia* to *D. simulans*, and preliminary analyses of interspecific introgression indicate that *D. simulans* and *D. sechellia* may currently be exchanging genes (Solignac and Monnerot 1986; Kliman et al. 2000; Garrigan et al. 2012; Brand et al. 2013). *Drosophila yakuba* and *D. santomea* hybridize in the midlands of Pico de São Tomé, and this hybrid zone has persisted since its discovery in 2000 (Lachaise et al. 2000). If we assume that *D. yakuba‘s* preference for open, agricultural areas has not changed over time, both the *D. yakuba-D. santomea* hybrid zone, and the *D. yakuba-D. teissieri* hybrid zone described here, may be relatively new. In the case of the *D. yakuba-D. teissieri* hybrid zone, the secondary forest where *D. teissieri* dwells is also recent, having grown after the downfall of the military dictatorship in Equatorial Guinea (1924-1979). Thus, while the exact dates are uncertain, agricultural expansion suggests that secondary contact in the *D. yakuba* clade may be recent.

The discovery of the *D. yakuba-D. teissieri* hybrid zone on Bioko (versus other areas of range overlap) may not be surprising, as hybridization is thought to be common in this region of Africa. For example, the reed frogs *Hyperolius molleri* and *H. thomensis* hybridize in the midlands of Pico de São Tomé in an area that directly overlaps with the *D. yakuba-D. santomea* hybrid zone (Bell et al. 2015). Limited data suggest that Caecilians *(Schistometopum thomense* and *S. ephele*) may also hybridize on São Tomé (Stoelting et al. 2014), Bioko populations of M and S *Anopheles gambiae* mosquitoes display evidence of past introgression (Lee et al. 2013), and hybridization may also occur in other systems (Melo 2006; Melo and O’Ryan 2007). An obvious question moving forward is whether hybridization is particularly common in this archipelago.

The distribution of phenotypic and genetic variation observed for hybrid individuals can provide an understanding of patterns of selection for or against hybrids. For example, Lindke et al. (2014) quantified the rate of hybridization between *Populus alba* and *P. tremula* in nature and showed that many of the recombinant genotypes present in germinated seed-sets are not recruited into the adult population. Selection acting on recombinant genotypes seems to contribute to RI between *P. alba* and *P. tremula* (Christe et al. 2016). More subtle patterns of genetic variation observed within advanced-generation hybrids, such as the overrepresentation of certain parental alleles in linkage disequilibrium can be used to infer how selection acts to maintain species boundaries (Schumer et al. 2014). Our results add to these studies by suggesting a phenotypic mechanism that likely contributes to a lack of advanced-generation hybrids in the Bioko *D. yakuba-D. teissieri* hybrid zone—namely,alternate patterns of dominance between parental alleles can result in a maladaptive combination of ecologically relevant phenotypes.

## conclusion

Areas where two species interbreed and hybridize provide unique windows into process and consequences of divergence. The study of secondary contact and hybrid zones has given rise to a full legacy of theoretical predictions and the development of empirical approaches to understand the dynamics of interspecific gene flow (Barton 1979; Barton and Hewitt 1985; Larson et al. 2014; Harrison and Larson 2016). The identification of hybrid zones in *Drosophila* has been rare and it is unclear how many areas of secondary contact exist in the genus. The Bioko *D. yakuba-D. teissieri* hybrid zone is exceptionally divergent and illustrates how, along with endogenous traits like hybrid male sterility, a maladaptive combination of ecologically-relevant parental phenotypes in hybrids may contribute to RI and maintain this seemingly stable, narrow, hybrid zone centered on the forest-open habitat ecotone. Discovering additional hybrid zones between *D. yakuba* and *D. teissieri*, across the whole African continent, would be useful to determine the consistency of selection against hybridization. However, identifying these hybrid zones might be a race against the clock because deforestation of *Parinari* trees is rapidly removing *D. teissieri’*s predicted native habitat (Cobb et al. 2000).

## Acknowledgements

We thank Matthew Hahn and Erica Larson for comments on an earlier draft that improved this paper, and Patrick Reilly and Peter Andolfatto for help with MSG. We also thank Patrick McLaughlin, Barret Milles, Gabriel Ousmane and Rayna Bell for their field assistance. Bioko Biodiversity Protection Program and the Universidad Nacional de Guinea Ecuatorial (UNGE) helped our research by kindly allowing us to use their facilities. Research reported in this publication was supported by startup funds to D.R.M. B.S.C. was supported by the National Institute of Allergy and Infectious Diseases of the National Institutes of Health (NIH) under Award Number F32AI114176. A.S. was supported by NIH R01 GM114093 to P. Andolfatto. The content is solely the responsibility of the authors and does not necessarily represent the official views of the NIH.

## Supplementary Material

**TABLE S1.** Preliminary releases of *D. melanogaster* subgroup species indicated that flies on average travel less than 200 m in 24 hr. *Drosophila yakuba* and *D. teissieri* travel under 100 m on average in 24 hr. The total number of flies released (*N*_REL_), the total number of flies recaptured (*N*_REC_), the mean distance (m) traveled from the release point, and standard deviations (SD) are reported.

| Species | $N_{REL}$ | $N_{REC}$ | Mean distance (m) | SD |
| --- | --- | --- | --- | --- |
| *D. yakuba* | 500 | 25 | 84.0 | 94.34 |
| *D. teissieri* | 500 | 22 | 45.45 | 67.10 |
| *D. melanogaster* | 500 | 16 | 93.75 | 99.79 |
| *D. simulans* | 500 | 33 | 190.91 | 112.82 |

**TABLE S2.** Desiccation tolerance of *D. yakuba* and *D. teissieri*, F_1_(♀*tei* × ♂*yak*), and F_1_(♀*yak* × ♂*tei*) genotypes by sex. Desiccation tolerance is measured as the mean number of hours a fly survives when exposed to desiccating conditions. Males are more prone to suffer from desiccation than are females (Cox hazards regression, sex effect: χ^2^= 8.39, df = 1, *P* = 0.004). The largest differences were observed between genotypes (Cox hazards regression, genotype effect: χ^2^ = 143.55, df = 3, *P*< 0.0001). Statistical analyses for pooled males and females are presented in Table 4.

|  | Female |  |  | Male |  |  |
| --- | --- | --- | --- | --- | --- | --- |
|  | *N* | Mean | SD | *N* | Mean | SD |
| *D. teissieri* | 99 | 4.84 | 1.76 | 99 | 4.62 | 1.61 |
| *D. yakuba* | 99 | 6.93 | 1.59 | 99 | 6.32 | 1.42 |
| F<sub>1</sub>(♀*tei* × ♂*yak*) | 99 | 4.78 | 1.38 | 99 | 4.74 | 1.61 |
| F<sub>1</sub>(♀*yak* × ♂*tei*) | 99 | 4.76 | 1.49 | 99 | 4.24 | 1.41 |

## Literature Cited

Adrion, J. R., M. W. Hahn, and B. S. Cooper. 2015. Revisiting classic clines in *Drosophila melanogaster* in the age of genomics. Trends in Genetics 31:434–444.

Andolfatto, P., D. Davison, D. Erezyilmaz, T. T. Hu, J. Mast, T. Sunayama-Morita, and D. L. Stern. 2011. Multiplexed shotgun genotyping for rapid and efficient genetic mapping. Genome Research 21:610–617.

Bachtrog, D., K. Thornton, A. Clark, and P. Andolfatto. 2006. Extensive introgression of mitochondrial DNA relative to nuclear genes in the *Drosophila yakuba* species group. Evolution 60:292–302.

Barton, N. H. 1979. Dynamics of hybrid zones. Evolution 43:341–359.

Barton, N. H. and G. M. Hewitt. 1985. Analysis of hybrid zones. Annual Review of Ecology and Systematics:113–148.

Bates, D., M. Mächler,B. Bolker, and S. Walker. 2015. Fitting Linear Mixed-Effects Models Using lme4. 2015 67:48.

Bell, R. C., R. C. Drewes, and K. R. Zamudio. 2015. Reed frog diversification in the Gulf of Guinea: Overseas dispersal, the progression rule, and *in situ* speciation. Evolution 69:904–915.

Brand, C. L., S. B. Kingan, L. Wu, and D. Garrigan. 2013. A Selective Sweep across Species Boundaries in *Drosophila*. Molecular Biology and Evolution 30:2177–2186.

Christe, C., K. N. Stölting, L. Bresadola,B. Fussi, B. Heinze, D. Wegmann, and C. Lexer. 2016. Selection against recombinant hybrids maintains reproductive isolation in hybridizing *Populus* species despite F1 fertility and recurrent gene flow. Molecular Ecology 25:2482–2498.

Cobb, M., M. Huet, D. Lachaise, and M. Veuille. 2000. Fragmented forests, evolving flies: molecular variation in African populations of *Drosophila teissieri*. Molecular Ecology 9:1591–1597.

Comeault, A. A., A. G. Serrato-Capuchina, D. A. Turissini, P. J. McLaughlin, J. R. David, and D. R. Matute. in press. A non-random subset of olfactory genes is associated with host preference in the fruit fly *Drosophila orena*. Evolution Letters.

Comeault, A. A., A. Venkat, and D. R. Matute. 2016. Correlated evolution of male and female reproductive traits drive a cascading effect of reinforcement in *Drosophila yakuba*. Proceedings of the Royal Society B: Biological Sciences 283.

Cooper, B. S., P. S. Ginsberg, M. Turelli, and D. R. Matute. 2017. *Wolbachia* in the *Drosophila yakuba* complex: pervasive frequency variation and weak cytoplasmic incompatibility, but no apparent effect on reproductive isolation. Genetics 205:333–351.

Cooper, B. S., L. A. Hammad, and K. L. Montooth. 2014. Thermal adaptation of cellular membanes in natural populations of *Drosophila melanogaster*. Functional Ecology 28:886–894.

Counterman, B. A. and M. A. F. Noor. 2006. Multilocus test for introgression between the cactophilic species *Drosophila mojavensis* and *Drosophila arizonae*. American Naturalist 168:682–696.

Coyne, J. A. and H. A. Orr. 1989. Patterns of speciation in *Drosophila*. Evolution 43:362–381.

Coyne, J. A. and H. A. Orr. 2004. Speciation. Sinauer Associates, Sunderland, MA.

Cruickshank, T. E. and M. W. Hahn. 2014. Reanalysis suggests that genomic islands of speciation are due to reduced diversity, not reduced gene flow. Molecular Ecology 23:3133–3157.

David, J. R., F. Lemeunier, L. Tsacas, and A. Yassin. 2007. The historical discovery of the nine species in the *Drosophila melanogaster* species subgroup. Genetics 177:1969–1973.

Dobzhansky, T. and T. G. Dobzhansky. 1937. Genetics and the Origin of Species. Columbia University Press.

Enjin, A., E. E. Zaharieva, D. D. Frank, S. Mansourian, G. S. Suh, M. Gallio, and M. C. Stensmyr. 2016. Humidity sensing in *Drosophila*. Current Biology : CB 26:1352–1358.

Funk, D. J., P. Nosil, and W. J. Etges. 2006. Ecological divergence exhibits consistently positive associations with reproductive isolation across disparate taxa. Proceedings of the National Academy of Sciences of the United States of America 103:3209–3213.

Garrigan, D., S. B. Kingan, A. J. Geneva, P. Andolfatto, A. G. Clark, K. R. Thornton, and D. C. Presgraves. 2012. Genome sequencing reveals complex speciation in the *Drosophila simulans* clade. Genome Research 22:1499–1511.

Grant, P. R. and R. Grant. 1992. Hybridization of bird species. Science 256:193.

Harrell Jr., F. E. 2013. Regression Modeling Strategies. R Package.

Harrison, R. G. 1990. Hybrid zones: windows on evolutionary process. Oxford Surveys in Evolutionary Biology 7:69–128.

Harrison, R. G. and E. L. Larson. 2016. Heterogeneous genome divergence, differential introgression, and the origin and structure of hybrid zones. Molecular Ecology 25:2454–2466.

Hewitt, G. M. 1988. Hybrid zones-natural laboratories for evolutionary studies. Trends in Ecology & Evolution 3:158–167.

Hey, J., Y. Chung, and A. Sethuraman. 2015. On the occurrence of false positives in tests of migration under an isolation-with-migration model. Molecular Ecology 24:5078–5083.

Hey, J. and R. Nielsen. 2004. Multilocus methods for estimating population sizes, migration rates and divergence time, with applications to the divergence of *Drosophila pseudoobscura* and *D. persimilis*. Genetics 167:747–760.

Hey, J. and R. Nielsen. 2007. Integration within the Felsenstein equation for improved Markov chain Monte Carlo methods in population genetics. Proceedings of the National Academy of Sciences of the United States of America 104:2785–2790.

Hoffmann, A. and V. Kellermann. 2006. Revisiting heritable variation and limits to species distribution: recent developments. Israel Journal of Ecology & Evolution 52:247–261.

Hoffmann, A. A. and A. R. Weeks. 2007. Climatic selection on genes and traits after a 100 year-old invasion: a critical look at the temperate-tropical clines in *Drosophila melanogaster* from eastern Australia. Genetica 129:133–147.

Holland, N. D., L. Z. Holland, and A. Heimberg. 2015. Hybrids between the Florida Amphioxus *(Branchiostoma floridae)* and the Bahamas Lancelet *(Asymmetron lucayanum)*: developmental morphology and chromosome Counts. The Biological Bulletin 228:13–24.

Hothorn, T., F. Bretz, and P. Westfall. 2008. Simultaneous Inference in General Parametric Models. Biometrical Journal 50:346–363.

Jaenike, J., K. A. Dyer, C. Cornish, and M. S. Minhas. 2006. Asymmetrical reinforcement and *Wolbachia* infection in *Drosophila*. Plos Biology 4:1852–1862.

Kay, M. and J. Wobbrock. 2016. ARTool: Aligned Rank Transform for Nonparametric Factorial ANOVAs.

Kellermann, V., V. Loeschcke, A. A. Hoffmann, T. N. Kristensen, C. Fløjgaard, J. R. David, J. C. Svenning, and J. Overgaard. 2012a. Phylogenetic constraints in key functional traits behind species climate nices: patterns of desiccation and cold resistance across 95 *Drosophila* species. Evolution 66:3377–3389.

Kellermann, V., J. Overgaard, A. A. Hoffmann, C. Flojgaard, J. C. Svenning, and V. Loeschcke. 2012b. Upper thermal limits of *Drosophila* are linked to species distributions and strongly constrained phylogenetically. Proceedings of the National Academy of Sciences of the United States of America 109:16228–16233.

Kellermann, V., B. van Heerwaarden,C. M. Sgrò, and A. A. Hoffmann. 2009. Fundamental evolutionary limits in ecological traits drive *Drosophila* species distributions. Science 325:1244–1246.

Key, K. 1968. The concept of stasipatric speciation. Systematic Biology 17:14–22.

Kliman, R. M., P. Andolfatto, J. A. Coyne, F. Depaulis, M. Kreitman, A. J. Berry, J. McCarter, J. Wakeley, and J. Hey. 2000. The population genetics of the origin and divergence of the *Drosophila simulans* complex species. Genetics 156:1913–1931.

Kulathinal, R. J., L. S. Stevison, and M. A. F. Noor. 2009. The genomics of speciation in *Drosophila:* diversity, divergence, and introgression estimated using low-coverage genome sequencing. PLoS Genetics 5:e1000550.

Lachaise, D., M. Harry, M. Solignac, F. Lemeunier, V. Benassi, and M. L. Cariou. 2000. Evolutionary novelties in islands: *Drosophila santomea*, a new *melanogaster* sister species from São Tomé. Proceedings of the Royal Society B: Biological Sciences 267:1487–1495.

Lachaise, D., F. Lemeunier, and M. Veuille. 1981. Clinal variations in male genitalia in *Drosophila teissieri*. American Naturalist 117:600–608.

Larson, E. L., T. A. White, C. L. Ross, and R. G. Harrison. 2014. Gene flow and the maintenance of species boundaries. Molecular Ecology 23:1668–1678.

Lee, Y., C. D. Marsden, L. C. Norris, T. C. Collier, B. J. Main,A. Fofana, A. J. Cornel, and G. Lanzaro. 2013. Spatiotemporal dynamics of gene flow and hybrid fitness between the M and S forms of the malaria mosquito, *Anopheles gambiae*. Proceedings of the National Academy of Sciences of the United States of America 110:19854–19859.

Lenth, R. 2016a. Least-Squares Means: The R Package lsmeans. Journal of Statistical Software 69:1–33.

Lenth, R. V. 2016b. Least-Squares Means: The R Package lsmeans. 2016 69:33.

Lindtke, D., Z. Gompert, C. Lexer, and C. A. Buerkle. 2014. Unexpected ancestry of *Populus* seedlings from a hybrid zone implies a large role for postzygotic selection in the maintenance of species. Molecular Ecology 23:4316–4330.

Llopart, A., D. Herrig, E. Brud, and Z. Stecklein. 2014. Sequential adaptive introgression of the mitochondrial genome in *Drosophila yakuba* and *Drosophila santomea*. Molecular Ecology 23:1124–1136.

Llopart, A., D. Lachaise, and J. A. Coyne. 2005a. An anomalous hybrid zone in *Drosophila*. Evolution 59:2602–2607.

Llopart, A., D. Lachaise, and J. A. Coyne. 2005b. Multilocus analysis of introgression between two sympatric sister species of *Drosophila: Drosophila yakuba* and *D. santomea*. Genetics 171:197–210.

Lohse, K., M. Clarke, M. G. Ritchie, and W. J. Etges. 2015. Genome - wide tests for introgression between cactophilic *Drosophila* implicate a role of inversions during speciation. Evolution 69:1178–1190.

Mahalanobis, P. C. 1936. On the generalised distance in statistics. Pp. 49–55. Proceedings National Institute of Science, India.

Maheshwari, S. and D. A. Barbash. 2011. The genetics of hybrid incompatibilities. Annual Review of Genetics 45:331–355.

Mallet, J. 2005. Hybridization as an invasion of the genome. Trends in Ecology & Evolution 20:229–237.

Mallet, J., N. Besansky, and M. W. Hahn. 2016. How reticulated are species? BioEssays 38:140–149.

Markow, T. and P. O’Grady. 2005. Drosophila: A Guide of Species Identification and Use.

Matute, D. R. 2010. Reinforcement of gametic isolation in *Drosophila*. PLoS Biology 8:e1000341.

Matute, D. R. and J. F. Ayroles. 2014. Hybridization occurs between *Drosophila simulans* and *sechellia* in the Seychelles archipelago. Journal of Evolutionary Biology 27:1057–1068.

Matute, D. R., I. A. Butler, D. A. Turissini, and J. A. Coyne. 2010. A test of the snowball theory for the rate of evolution of hybrid incompatibilities. Science 329:1518–1521.

Matute, D. R. and A. Harris. 2013. The influence of abdominal pigmentation on desiccation and ultraviolet resistance in two species of *Drosophila*. Evolution 67:2451–2460.

McKenna, A., M. Hanna, E. Banks, A. Sivachenko, K. Cibulskis, A. Kernytsky, K. Garimella, D. Altshuler, S. Gabriel, M. Daly, and M. A. DePristo. 2010. The Genome Analysis Toolkit: a MapReduce framework for analyzing next-generation DNA sequencing data. Genome Research 20:1297–1303.

Melo, M. 2006. Bird speciation in the Gulf of Guinea. University of Edinburgh, Edinburgh.

Melo, M. and C. O’Ryan. 2007. Genetic differentiation between Príncipe Island and mainland populations of the grey parrot *(Psittacus erithacus)*, and implications for conservation. Molecular Ecology 16:1673–1685.

Mendelson, T. C. 2003. Sexual isolation evolves faster than hybrid inviability in diverse sexually dimorphic genus of fish (Percidae: *Etheostoma*). Evolution 57:317–327.

Monnerot, M., M. Solignac, and D. R. Wolstenholme. 1990. Discrepancy in divergence of the mitochondrial and nuclear genomes of *Drosophila teissieri* and *Drosophila yakuba*. Journal of Molecular Evolution 30:500–508.

Moyle, L. C., M. S. Olson, and P. Tiffin. 2004. Patterns of reproductive isolation in three angiosperm genera. Evolution 58:1195–1208.

Muller, H. J. 1942. Isolating mechanisms, evolution and temperature. Pp. 71–125. Biol. Symp.

Picelli, S., Å. K. Björklund, B. Reinius, S. Sagasser, G. Winberg, and R. Sandberg. 2014. Tn5 transposase and tagmentation procedures for massively scaled sequencing projects. Genome Research 24:2033–2040.

Price, T. D. and M. M. Bouvier. 2002. The evolution of F1 postzygotic incompatibilities in birds. Evolution 56:2083–2089.

Rio, B., G. Couturier, F. Lemeunier, and D. Lachaise. 1983. Evolution d’une specialisation saisonniere chez *Drosophila erecta* (Dipt., Drosophilidae). Annales de la Société Entomologique de France 2:235–248.

Rothfels, C. J., A. K. Johnson, P. H. Hovenkamp, D. L. Swofford, H. C. Roskam, C. R. Fraser-Jenkins, M. D. Windham, and K. M. Pryer. 2015. Natural hybridization between genera that diverged from each other approximately 60 million years ago. American Naturalist 185:433–442.

Schumer, M., R. Cui, D. L. Powell, R. Dresner, G. G. Rosenthal, and P. Andolfatto. 2014. Highresolution mapping reveals hundreds of genetic incompatibilities in hybridizing fish species. eLife 3:e02535.

Shoemaker, D. D., V. Katju, and J. Jaenike. 1999. *Wolbachia* and the evolution of reproductive isolation between *Drosophilla recens* and *Drosophila subquinaria*. Evolution 53:1157–1164.

Singmann, H., B. Bolker, and J. Westfall. 2015. afex: Analysis of Factorial Experiments. R package version 0.13–145.

Smit, A., R. Hubley, and P. Green. 2013-2015. RepeatMasker 0pen–4.0.

Solignac, M. and M. Monnerot. 1986. Race formation, speciation, and introgression within *Drosophila simulans*, *D. mauritiana*, and *D. sechellia* inferred from mitochondrial DNA analysis. Evolution 40:531–539.

Stebbins, G. L. 1959. The role of hybridization in evolution. Proceedings of the American Philosophical Society 103:231–251.

Stoelting, R. E., G. J. Measey, and R. C. Drewes. 2014. Population Genetics of the São Tomé Caecilian (Gymnophiona: Dermophiidae: *Schistometopum thomense)* Reveals Strong Geographic Structuring. PLOS ONE 9:e104628.

Stukenbrock, E. H., F. B. Christiansen, T. T. Hansen, J. Y. Dutheil, and M. H. Schierup. 2012. Fusion of two divergent fungal individuals led to the recent emergence of a unique widespread pathogen species. Proceedings of the National Academy of Sciences of the United States of America 109:10954–10959.

Turelli, M., N. H. Barton, and J. A. Coyne. 2001. Theory and speciation. Trends in Ecology & Evolution 16:330–343.

Turelli, M., J. R. Lipkowitz, and Y. Brandvain. 2014. On the Coyne and Orr-igin of species:effects of intrinsic postzygotic isolation, ecological differentiation, *X* chromosome size, and sympatry on *Drosophila* speciation. Evolution 68:1176–1187.

Turelli, M. and L. C. Moyle. 2007. Asymmetric postmating isolation: Darwin’s corollary to Haldane’s rule. Genetics 176:1059–1088.

Turissini, D. A., A. A. Comeault, G. Liu, Y. C. G. Lee, and D. R. Matute. 2017. The ability of *Drosophila* hybrids to locate food declines with parental divergence. Evolution, 71: 960–973.

Turissini, D. A., G. Liu, J. R. David, and D. R. Matute. 2015. The evolution of reproductive isolation in the *Drosophila yakuba* complex of species. Journal of Evolutionary Biology. 28: 557–75

Turissini, D. A. and D. R. Matute. in revision. Fine scale mapping of genomic introgressions within the *Drosophila yakuba* clade.

White, G. C., D. R. Anderson, K. P. Burnham, and D. L. Otis. 1982. Capture-recapture and removal methods for sampling closed populations. Pp. 14, Los Alamos, NM.

Widmer, A., C. Lexer, and S. Cozzolino. 2009. Evolution of reproductive isolation in plants. Heredity (Edinb) 102:31–38.

Yassin, A. 2017. *Drosophila yakuba mayottensis*, a new model for the study of incipient ecological speciation. Fly 11:37–45.

Yassin, A., V. Debat, H. Bastide, N. Gidaszewski, J. R. David, and J. E. Pool. 2016. Recurrent specialization on a toxic fruit in an island *Drosophila* population. Proceedings of the National Academy of Sciences of the United States of America 113:4771–4776.

